# Deep visual multi-omics profiling reveals mechanisms that underlie cancer cell differentiation and aggressiveness in clear cell renal cell carcinoma

**DOI:** 10.1101/2025.01.31.635927

**Authors:** Hella A. Bolck, Ede Migh, Andras Kriston, Natalia Zajac, Susanne Kreutzer, Tiberiu Totu, Peter Leary, Ferenc Kovacs, Dorothea Rutishauser, Sibylle Pfammatter, Jonas Grossmann, Cassandra Litchfield, Marija Buljan, Niels J. Rupp, Peter Horvath, Holger Moch

## Abstract

Clear cell renal cell carcinoma (ccRCC) exhibits significant intra-tumoral heterogeneity (ITH) at both morphological and genetic levels, complicating treatment and contributing to disease progression. Among these, ccRCCs with focal rhabdoid differentiation stand out as highly aggressive tumors distinguished by cells with unique morphological features. However, the correlation between distinct morphological phenotypes, specific molecular alterations, and their influence on tumor behavior remains poorly understood.

In this study, we integrated advanced AI-based image analysis with single-cell isolation and multi-omics profiling to dissect the link between clinically relevant morphological and molecular features of ccRCC cells. Using a novel digital pathology workflow, we quantified low-grade, high-grade, and rhabdoid morphologies in ccRCC diagnostic images with unprecedented precision. Subsequently, isolation of two sets of 1,000 morphologically distinct cells for detailed mRNA and protein expression analyses, revealed significant increasing dysregulation associating with higher histopathological grades. Rhabdoid ccRCC cells (grade 4) demonstrated unique molecular profiles, including upregulated FOXM1-driven proliferation, disrupted cell-matrix interactions, and enhanced immune evasion pathways. Despite high T-cell infiltration in rhabdoid areas, we identified a rhabdoid-specific immunosuppressive network driven by cytokines, IFN-beta, and integrin signaling, likely contributing to T-cell exhaustion. Rhabdoid ccRCC cells develop a distinct immunosuppressive signaling network, involving PD-L1 and novel immunomodulatory factors such as CD38 and ITGB2. These findings provide a basis for novel therapeutic strategies targeting these pathways in combination with immunotherapy to improve outcomes for patients with aggressive rhabdoid ccRCC.

**Key Points:**

- ccRCC is characterized by well-established morphological heterogeneity but the correlation with the underlying molecular aberrations remained elusive.
- By integrating AI-based image analysis with single cell isolation and deep multi-omics profiling, we dissect the molecular intricacies of ccRCC, from targeted collection of 1,000 morphologically distinct cells.
- Our results demonstrate significant dysregulation of gene and protein expression correlating with higher histopathological grades in ccRCC.
- Aggressive ccRCC cells with rhabdoid differentiation (grade 4) display distinct molecular profiles, as they upregulate FOXM1-mediated proliferation, ECM remodeling and the immune evasion responses, suggesting new therapeutic avenues enhancing ICI efficacy in these patients.

## Introduction

Celar cell renal cell carcinoma (ccRCC) is the most common subtype of renal cell carcinoma (RCC), accounting for the majority of metastatic cases and RCC-related deaths^1–3^. The hallmark genomic alteration in ccRCC is loss of chromosome 3p concurrent with biallelic inactivation of the *VHL* tumor suppressor gene. *VHL* loss drives constitutive activation of hypoxia-inducible pathways and profoundly alters cellular programs controlling angiogenesis, metabolism, and survival^4^. The early events also spur clonal evolution and profound genetic intra-tumor heterogeneity (ITH), a defining feature of ccRCC that complicates both diagnosis and treatment^5–7^.

Clinically, the WHO/ISUP grading system remains a cornerstone of prognosis and treatment guidance, stratifying ccRCCs into four grades, primarily based on nucleolar prominence^8,9^. Grade 4 tumors are additionally defined by the presence of sarcomatoid or rhabdoid differentiation, morphological features strongly associated with poor prognosis^9^. Yet ccRCCs often harbor a mosaic of low- (grade 1/2) and high-grade (grade 3/4) phenotypes^10^, highlighting a striking histo-morphological ITH^11^. However, the molecular underpinnings of this morphological heterogeneity, particularly in grade 4 tumors, remain elusive.

Recently, sarcomatoid RCCs have received growing attention^12,13^ as they have shown shown notable responses to immune checkpoint inhibitors (ICI) treatments^13–16^ which is potentially linked to exhibiting increased tumor-infiltrating lymphocytes (TILs)^1,17^. Much less is known about the molecular landscape and treatment outcomes for ccRCC with rhabdoid dedifferentiation^12^, that are observed in 3-7% of RCC cases^18–20^. The available post-hoc data suggests that these may also benefit from ICI treatment^21^ similarly to sarcomatoid cases. However, the mechanisms behind their putative ICI response remains largely unknown. Unravelling the molecular drivers of rhabdoid morphology and its tumor–immune interactions could unveil vulnerabilities in these aggressive tumors.

Recent advances in artificial intelligence (AI)-based digital pathology, single-cell RNA sequencing (RNAseq), and ultra-sensitive proteomics have created new opportunities to dissect tumor heterogeneity at cellular resolution *in situ*^22–25^. However, these technologies have yet to be combined into a single framework for linking tumor morphology to the molecular states to advance our understanding of cell-to-cell variation within tumors^22^.

Here, we present Deep Visual Multi-Omics, an integrated transformative workflow that bridges AI-guided histological cell classification, single-cell laser-capture microdissection, and ultra-sensitive multi-omics data collection from FFPE tissue samples. Applied to clinical ccRCC samples, we map gene and protein expression programs across thousands of morphologically defined cells providing biological insights into molecular programs that drive cellular differentiation and define morphological ITH. Our findings provide mechanistic insights for understanding the response of rhabdoid ccRCC to ICIs and offer new therapeutic strategies targeting these most aggressive subsets of tumor cells.

## Results

### Genome-wide genetic profiling of five ccRCCs with pronounced histo-morphological intra-tumor heterogeneity (ITH)

We selected five ccRCC cases exhibiting pronounced tumor-intrinsic micro-heterogeneity defined by different grade morphologies within a tumor (Fig. 1A and Suppl. Fig. 1A, Suppl. Table 1). Each sample underwent expert pathological assessment, confirming the co-occurrence of grade 1/2 (low grade), grade 3 (high grade), and rhabdoid (grade 4) ccRCC areas. Subsequently, we dissected gross tumor areas from all morphologies and adjacent normal tissue (Suppl. Fig. 1B) and subjected them to whole-genome DNA sequencing (WGS) (Suppl. Fig. 1C and 1D). Our analysis revealed that the tumor mutational burden (TMB) remained consistent across samples aligning with previously reported values in ccRCC with and without rhabdoid features^6,13^ (Fig. 1B). Arm-level chromosomal aberrations included the most prevalent structural alterations found in ccRCC, 3p loss and 5q gain (Fig. 1B and Suppl. Fig. 1E)^7^. Additionally, we consistently identified loss of chromosome 14q, a copy number variation (CNV) that has been associated with aggressive disease ^26,27^. Our findings also include previously reported CNVs, such as gains in chromosomes 7q and 9p, as well as loss of chromosome 8q ^27–29^. Notably, all ccRCC samples exhibited mutations in the *VHL* tumor suppressor gene (N=5) and harbored additional mutations in SETD2 (N=4), PBRM1 (N=4), BAP1 (N=2) as well as other frequently mutated ccRCC drivers (Fig. 1B, Suppl. Table 2 and 3)^6,27^. Thus, our genomic characterization largely corroborated findings from prior studies, providing confirmation of the typical molecular profiles of ccRCC samples.

**Figure 1:**
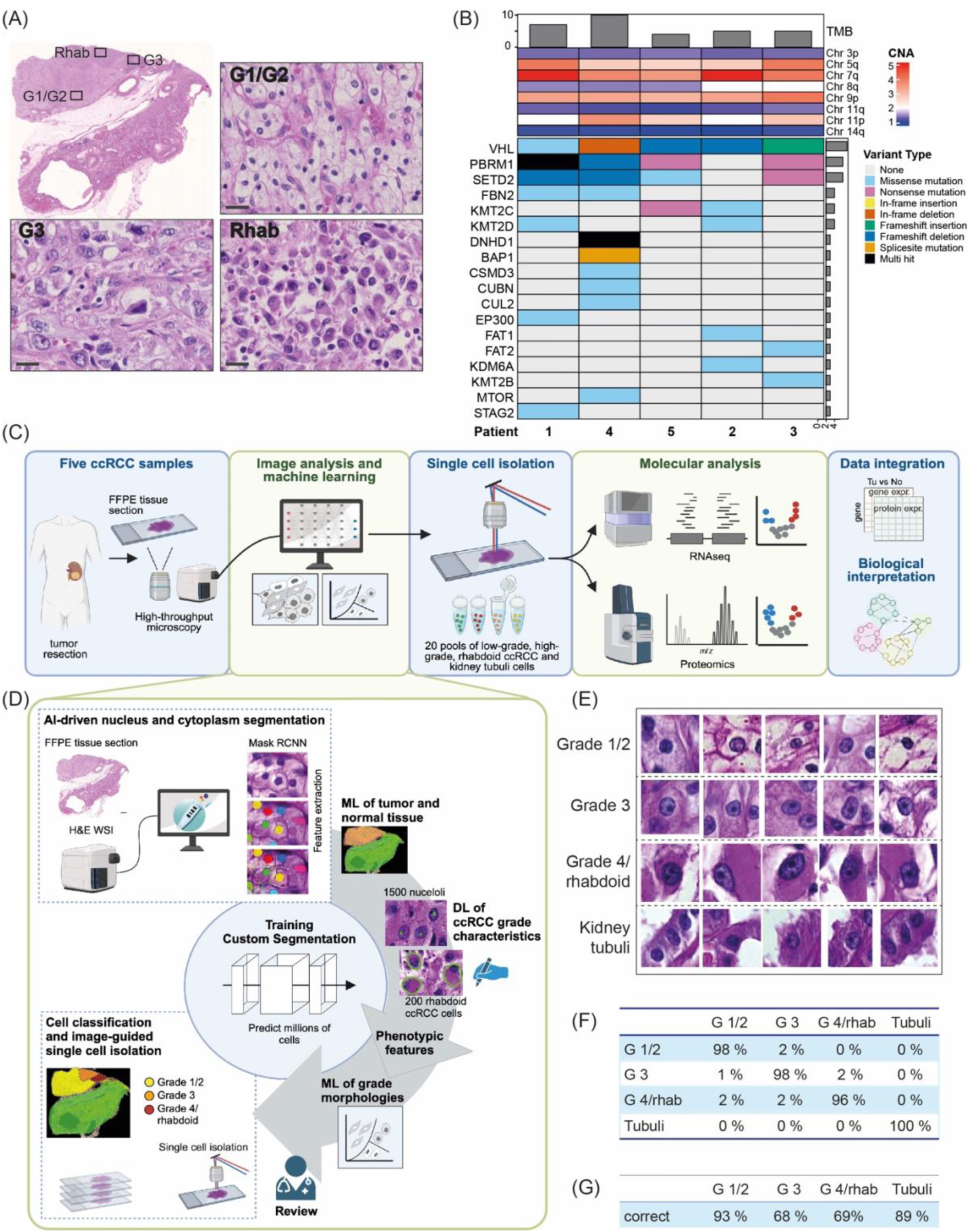
AI-driven, image-guided single cell isolation and multi-omics analysis of ccRCC with pronounced morphological intra-tumor heterogeneity including rhabdoid de-differentiation. (A) Representative H&E images (Patient 4) indicating ITH of grade features: Grade 1/2 (low-grade, G1/2), grade 3 (high-grade, G3) and grade 4 ccRCC with rhabdoid differentiation (Rhab). Scale bars: WSI – 2 mm, high magnification panels - 100 μm. (B) Overview of somatic alterations identified by WGS analysis in five samples used in this study, including TMB, selected copy number events and driver mutations. (C) Overview of workflow combining AI-guided single-cell histological classification and isolation with transcriptomic and proteomic analysis. (D) AI-driven segmentation workflow for grade 1/2 (low), grade 3 (high) and grade 4/rhabdoid ccRCC as well as normal-appearing kidney tubuli cells in diagnostic tissue image using BIAS. (E) Example images of cells from the classes stated above identified by ML. (F) Cross-validation accuracies of our image analysis workflow using a randomized 10-fold validation test. (G) Validation using independent unseen data.

### A deep digital pathology workflow quantified morphological heterogeneity in diagnostic images

Next, we used artificial intelligence-based image analysis on the HE-stained diagnostic whole-slide images (WSIs) of these cases to quantify morphological heterogeneity automatically (Fig. 1C and 1D). This involved the BIAS software suite^30^ for deep learning-based segmentation and phenotype classification. A pre-trained neural network segmented nuclei and subsequently features from both nuclei and surrounding areas were extracted^31^. An artificial neural network (ANN) classified regions as ccRCC or normal kidney tissue. A Mask R convolutional neural network (R-CNN) was trained by manual annotation of more than 1200 nucleoli to recognize grade 3 ccRCC cells (Suppl. Table 4). Similarly, we developed a model to segment rhabdoid differentiation in ccRCC cells (Grade 4) using a fine-tuned Mask R CNN that was trained by annotating more than 250 rhabdoid ccRCC cells (Suppl. Table 4). The results of these DL-based segmentation algorithms were integrated as additional features of putative tumor cells to train a classifier distinguishing grade 1/2 (low-grade), grade 3 (high-grade), and grade 4 (rhabdoid) ccRCC cells (Fig. 1D and 1E). Subsequently, we predicted these classes in two to three WSIs from each patient in our cohort and validated the model using a 10-fold self-validation test and on an unseen dataset in comparison to pathologist-annotated ground truth (Fig. 1F and 1G). Applying out workflow, we subsequently selected two sets of 1000 cells confidently classified as either grade 1/2 (low-grade), grade 3 (high-grade), grade 4 ccRCC cells with rhabdoid differentiation or normal kidney tubuli for each of the five ccRCC patient tissues and performed single-cell isolation (Fig. 1C).

### mRNA and protein-level alterations in ccRCC cells with morphologic heterogeneity reflect core disease genes and pathways

Following cell isolation, one replicate pool per morphological class was subjected to RNAseq analysis employing a protocol tailored for ultra-low sample inputs, which directly utilizes total RNA for library preparation input and relies on random priming. As a result, we obtained RNA expression data that mapped to 20,528 genes (Fig. 2A and Suppl. Table 5). The normal tubuli cells for patient 4 were found to display substantial tubular atrophy and interstitial fibrosis and were therefore excluded from further analysis. Principle component analysis (PCA) and hierarchical clustering analysis of the gene expression data revealed a clear distinction between the profiles of adjacent normal tubuli and ccRCC cells of various morphologies (Fig. 2A and Suppl. Fig. 2A). The PCA distances between ccRCC histologies were notably lower aligning with the expected similarity between these samples. However, a discernible trend of separation emerged between low- and higher-grade morphologies, particularly along PC1. Subsequently, we compared each of the ccRCC grade groups to the adjacent normal cells to determine differentially expressed genes (DEGs). The analysis revealed a significant increase of the number of both down- and upregulated mRNAs with higher grades (FDR <0.05 and |Log2FC| > 1) (Fig. 2B). Notably, the number of deregulated genes exhibited an over 4-fold increase between grade 1/2 ccRCC cells and the highest grade 4 rhabdoid cells, indicating a progressive dysregulation pattern.

**Figure 2:**
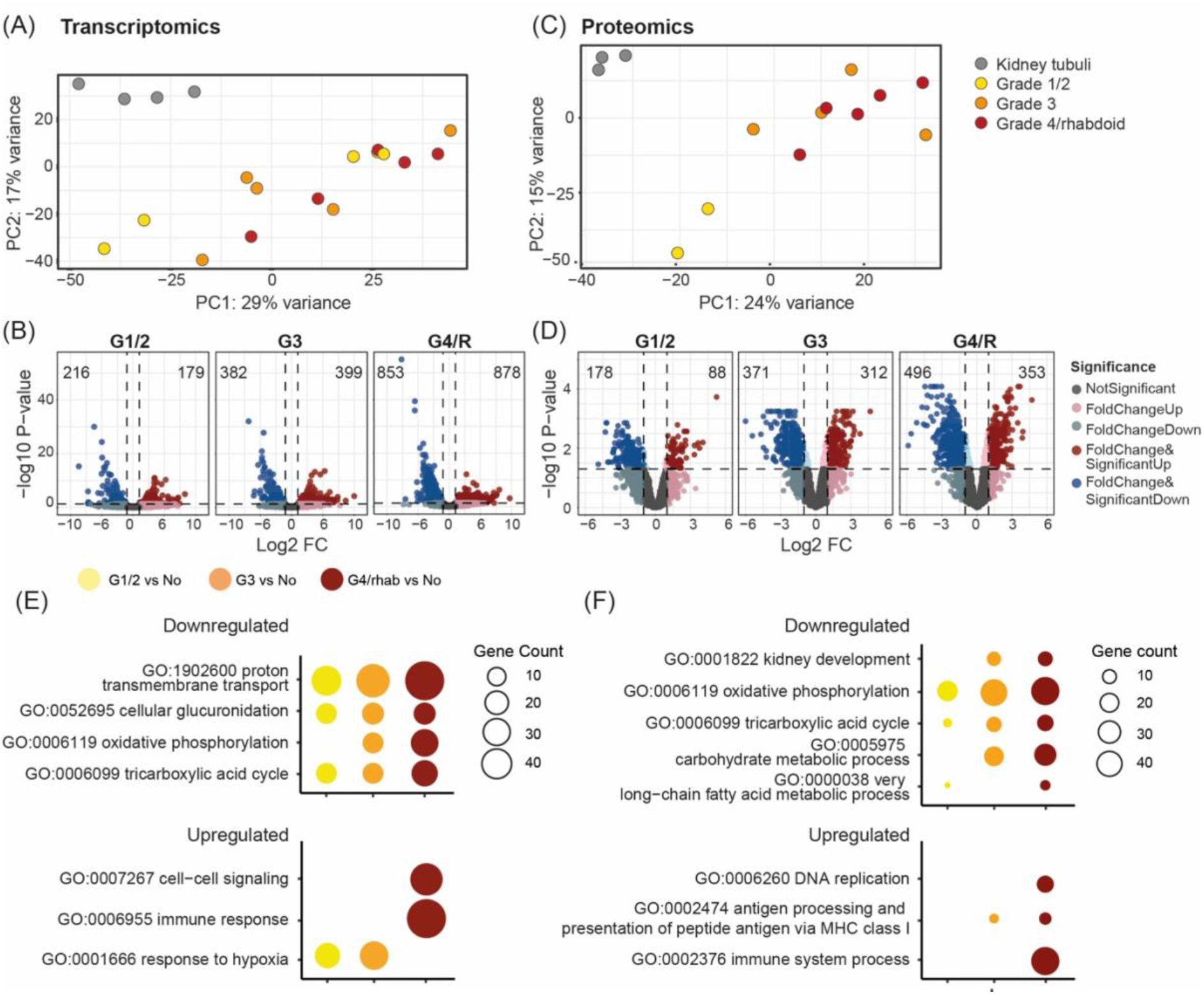
Differential Expression Analysis of mRNAs and proteins from individually isolated cells with distinct grade morphologies. (A) PCA of RNAseq data for 20,528 genes from five patients. (B) Volcano plot showing DEGs (FDR < 0.05, logFC > |1|) in ccRCC grade groups compared to adjacent normal tissues. Genes that were significantly down- or upregulated are presented with red/blue filled scatters and their number is indicated in the upper left and right corners, respectively. (C) PCA of LC-MS data for 4,382 proteins from five patients. (D) Volcano plot showing DEPs (FDR < 0.05, logFC > |1|) in ccRCC grade groups compared to adjacent normal tissues. Proteins that were significantly down- or upregulated are presented with red/blue filled scatters and their number is indicated in the upper left and right corners, respectively. (E) Enrichment analysis of associated GO biological pathways generated from over-representation analysis (hypergeometric test) for the transcriptomic datasets indicating adjusted p-values. (F) Same as (E) for proteomic dataset.

Next, we investigated proteome alterations in ccRCC cells with heterogeneous histologies by subjecting the second pool of isolated cells to LC-MS adapting a highly sensitivity workflow compatible with ultra-low sample inputs^22^. Employing data-independent acquisition with parallel accumulation–serial fragmentation (diaPASEF) with an additional ion mobility dimension (TIMS), we successfully identified 4,382 proteins. Six samples were excluded from further analysis due to irregularities during laser microdissection (LMD) that resulted in lower protein counts and noticeable batch effects (Suppl. Table 5). PCA and hierarchical clustering analysis of the remaining proteomics data confirmed a distinct separation of the normal tubuli and all morphological classes of ccRCC cells (Fig. 2C and Suppl. Fig. 2B). Notably, the gradual transition among the ccRCC samples with increasing grades, as expressed by PCA distances, was more pronounced than observed in the transcriptomic data, particularly between cells with low (Grade 1/2) and higher (Grade 3/4) grades (Fig. 2C). Moreover, differential expression analysis between ccRCC grade morphologies and their adjacent normal tubuli cells revealed an increasing number of differentially expressed proteins (DEPs) correlating with higher ccRCC grade, aligning with the trend observed in the transcriptomic data (Fig. 2D).

Overrepresentation analysis of the respective datasets revealed that downregulation of biological processes was more prominent than upregulation. Biological enrichment in DEG and DEP downregulated datasets uncovered terms mirroring known ccRCC-specific tumorigenic changes including oxidative phosphorylation and tricarboxylic acid cycle (TCA). Similarly, pathways related to normal kidney function (kidney development, glucuronidation, proton transmembrane transport) were increasingly downregulated with higher ccRCC grades (Fig. 2E and 2F, Suppl. Table 6 and 7). In contrast, biological pathways enriched in the upregulated datasets included response to hypoxia, DNA replication and cell-cell signaling. Notably, among the pathways upregulated specifically in grade 4/rhabdoid cells, we identified the immune response and related processes.

Taken together, our data established that AI-driven automated classification of ccRCC cells with distinct histological features followed by their individual LMD from formalin-fixed paraffin-embedded (FFPE) tissue from the classical diagnostic workup yielded suitable samples for deep molecular profiling.

### Functional overlap between mRNA and protein dysregulation reveals progressive tumorigenic changes associate with increasing tumor grades

Even though regulation of transcription and protein expression can be distinct, we observed notable correlation between mRNA and protein differential expression in our data (Fig. 3A). We detected 40, 134 and 245 significant genes shared across the mRNA and protein data for grade 1/2, grade 3 and grade 4/rhabdoid cells, respectively (Suppl. Fig. 3A). Importantly, Vimentin (VIM) a key diagnostic marker indicating epithelial-to-mesenchymal transition (EMT) was found strongly elevated in both datasets, while other well-established ccRCC markers such as CA9 and CAV1, showed strong positivity in at least one of the -omics layers (Fig. 3A). Despite their overlap, the majority of regulated genes were distinct to one of the two data layers (Fig. 3A and Suppl. Fig. 3A) prompting us to investigate whether the regulated genes drive similar biological functions (Fig. 3B and Suppl. Fig. 3B). Enrichment analyses uncovered widespread metabolic reprogramming in glucose and lipid metabolism as well as the repression of pathways related to physiological kidney functions in both datasets. Interestingly, often these alterations were observed across all ccRCC grades (Suppl. Fig. 3B). However, the number of overlapping biological pathways increased concomitantly with tumor grade, reinforcing the link between dysregulation and higher-grade morphologies^4^. The only upregulated biological pathway that was shared by mRNA and protein datasets and distinct to ccRCC cells with rhabdoid differentiation was the "immune response" suggesting a crucial role for the transformation and aggressiveness of ccRCC, Consistent with the almost universal loss of VHL in ccRCC, upregulation of HIF-1 signaling was evident across all grade groups at both the mRNA and protein levels and HIF1α transcriptional targets, such as VEGFA and GLUT1 (SLC2A1), showed significant increases in both -omics layers. (Fig. 3C). Concomitantly, HIF-signaling-mediated metabolic adaptations were evident from the data indicated by elevated expression of glycolytic enzymes across all ccRCC morphologies (e.g., ENO2, PFKB, PKM) and repression of oxidative phosphorylation, signified by downregulation of electron transport chain enzymes (NDUFS1, NDUFV1 – Complex I; COX5A, COX6C – Complex IV; ATP6V1A, ATP6VB1 – Complex V) and TCA cycle components (ACO2, SDHA) (Fig. 3C). Moreover, we observed significant alterations in lipid metabolism, with fatty acid beta-oxidation enzymes (ACAT1, ECHS1, EHHADH) being among the most significantly downregulated processes in both datasets. We also found evidence for progressive dysregulation of the one-carbon metabolism, a crucial pathway for tumor progression that has been recently described in relation to ccRCC aggressiveness (Suppl. Fig. 3C)^32^. Reflecting the progressive loss of cellular identity during tumorigenesis and morphological transition into higher-grade components, we observed a gradual downregulation of proximal renal tubule genes and enzymes related to physiological kidney functions, such as glucuronidation and transmembrane transport, which correlated with higher tumor grades (Suppl. Fig. 3C). Furthermore, grade 3 and grade 4/rhabdoid cells exhibited significant upregulation of essential stem cell markers such as CD44 and DCLK1^33,34^.

**Figure 3:**
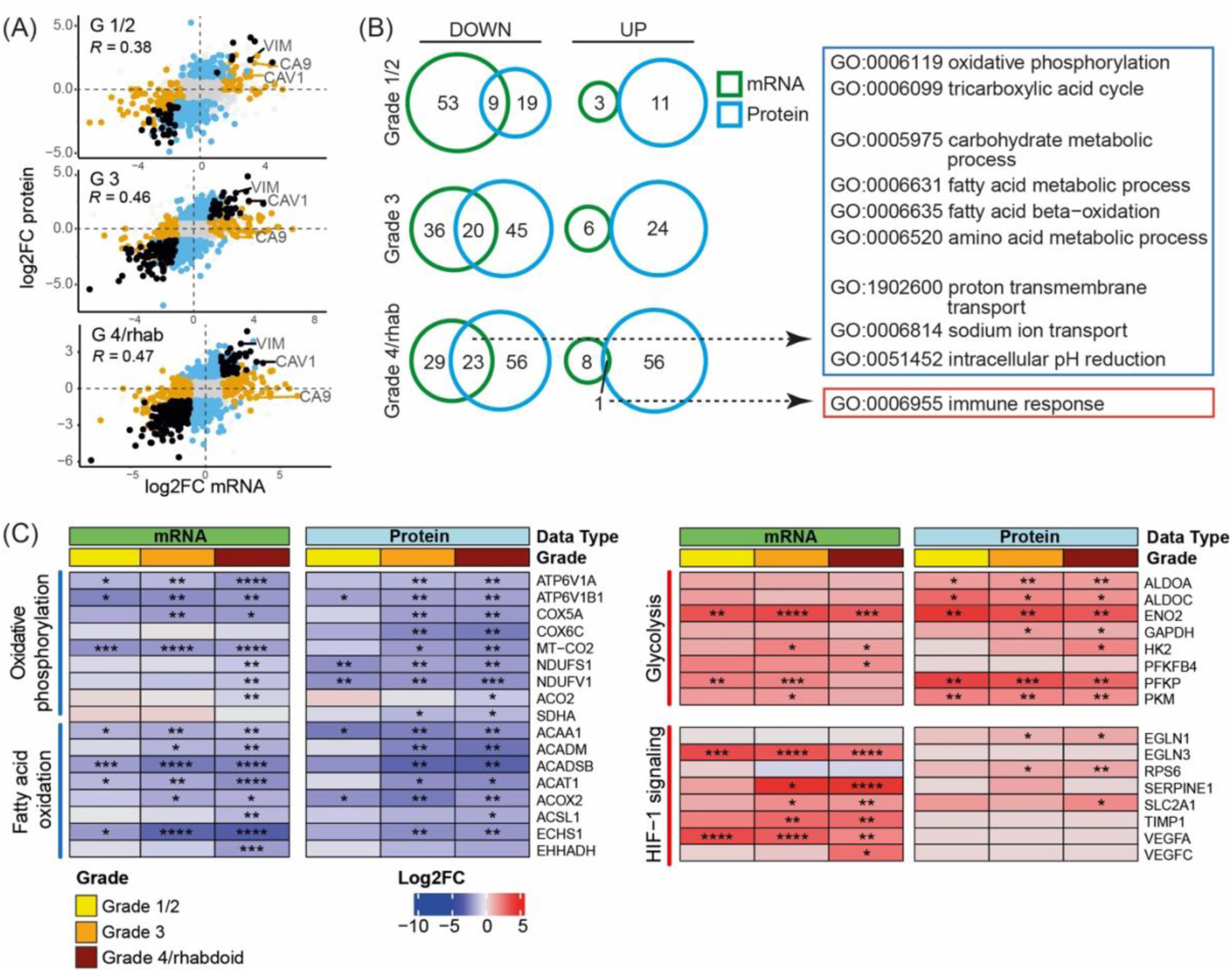
Functional overlap between mRNA and protein data. (A) Scatterplot of differential mRNA and protein expression between ccRCC grades and adjacent normal cells (Log2FC) indicating the correlation between datasets. Each dot represents one gene. Negative values indicate a decrease and positive values an increase in expression between tumor and NAT (B) Venn diagram showing numbers of enriched GO biological pathways in DEGs and DEPs for each grade group and the overlap between both data layers. Selected shared GO terms for grade 4/rhabdoid class of ccRCC cells are specified (for a complete overview, refer to Suppl. Fig. 4B). (C) Heatmaps depicting metabolic rewiring and upregulation of HIF1-signaling during ccRCC de-differentiation evident on both data layers. Alterations of representative genes and proteins are depicted as log2FC with respect to adjacent normal cells. Diagram on the left side depicts downregulated factors (blue), while on the right side upregulated factors are shown (red). Genes were selected from Glycolysis (Reactome R-HSA-70171), Fatty acid metabolism (Reactome R-HSA-8978868), Oxidative phosphorylation (KEGG path:hsa00190) and HIF-1 signaling (KEGG path:hsa04066). FDR is indicated by"****" <= 0.0001, "***" <= 0.001, "**"<= 0.01, "*" <= 0.05.

### Integrated network analysis unveils cell-matrix interactions and enhanced proliferation are key alterations characterizing rhabdoid ccRCC cells

As the direct overlap between mRNA and protein expression was limited (Fig. 3B), we employed integrative network-based analysis of transcriptomic and proteomic profiles to assess whether dysregulation on both layers would point to similar deregulated processes, offering deeper insights into the mechanisms underlying ccRCC tumor progression and histological heterogeneity. Specifically, we combined differentially expressed elements for both data layers and constructed grade-specific networks based on knowledge-based high confidence protein-protein interactions (PPI) networks. After constructing the networks, we used the current flow betweenness centrality metric in order to identify central nodes that connect elements in each network^35^. The top 30% most central elements of each network were further examined (Fig. 4A and Suppl. Table 10). Specifically, in the grade 4/rhabdoid-specific network, downregulated central nodes included factors implicated in renal functions (e.g. ACE2, NHERF1, ALDH3A2) as well as cell adhesion and EMT (e.g. CDH1). Conversely, in the network of upregulated factors, central elements reflected cellular adaptations related to processes such as EMT, cell migration, adhesion (e.g., VIM, ANXA2), and the Warburg effect (e.g., SLC2A1). Thus, the central network nodes that integrated contributions of mRNA and protein dysregulations corroborated the reprogramming of cellular processes towards a more aggressive and metastatic phenotype in grade 4/rhabdoid cells. We next applied MONET, a network modularization tool, which was constructed based on top performing algorithms for identifying modules (i.e. communities or subnetworks) of disease-related genes in the networks. We subsequently uncovered functionally related network modules associating with specific biological processes that could describe distinct changes in ccRCC cells with different grade morphologies (Fig. 4B). In Grade 4/rhabdoid ccRCC cells we determined a number of unique modules with biological functions not shared by the other morphologies (Fig. 4B and Suppl. Fig. 4A). Downregulated modules contained largely metabolic processes previously associated with ccRCC tumorigenesis underscoring that progressive transcriptional and translational dysregulation correlated with higher tumor grades.

**Figure 4:**
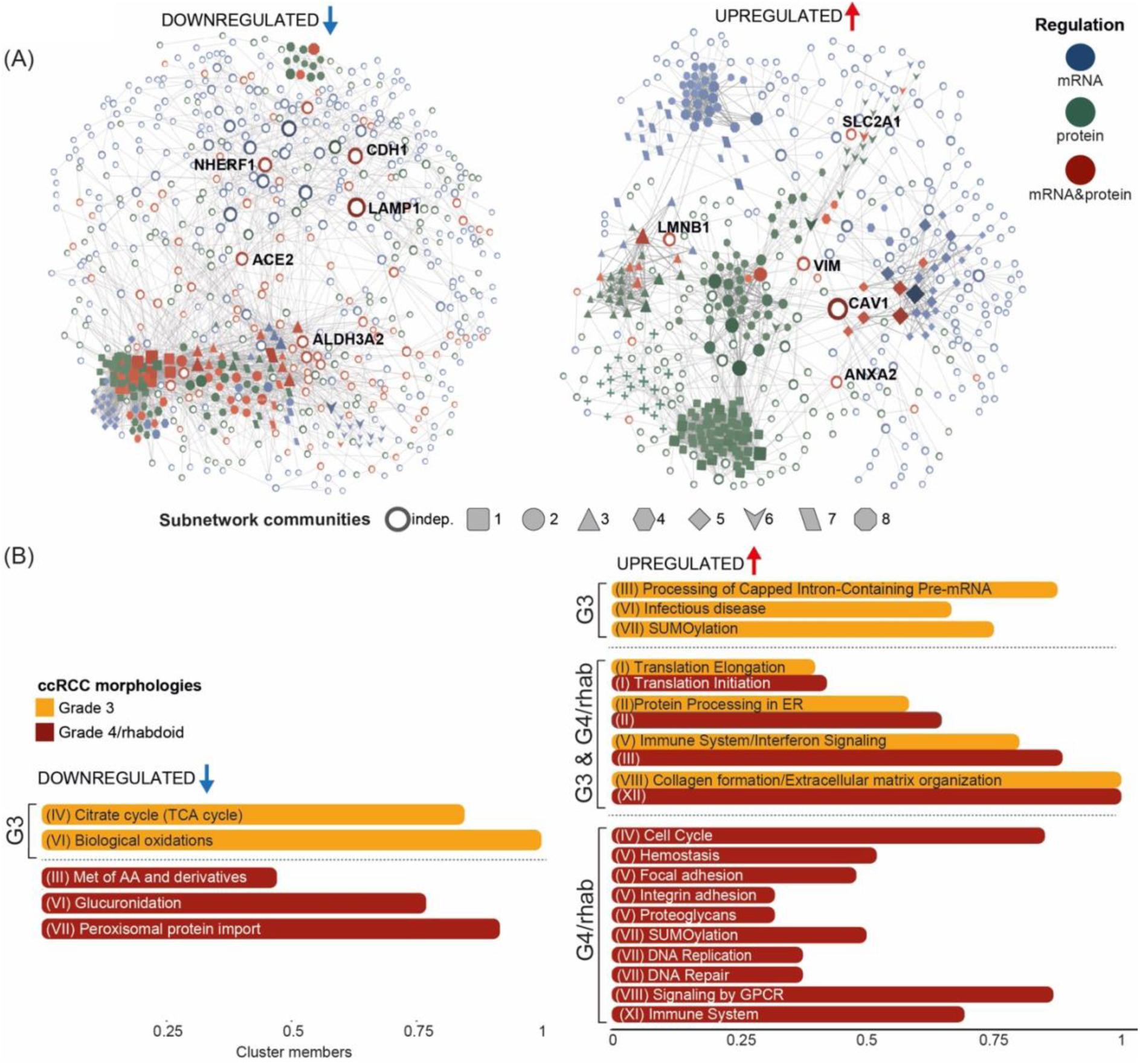
Multi-omics data integration uncovers distinct de-regulated processes in aggressive high-grade and rhabdoid ccRCC cells. (A) PPI network distinct to G4/rhabdoid ccRCC cells built by integrating the significant elements from transcriptomic and proteomic datasets that were down- (N=706) or upregulated (N=501). Node sizes and darker colors correspond to higher centrality scores. The five top highly connected hub genes that associated with no specific subnetwork are highlighted. Elements within network communities with defined biological functions, identified after the network decomposition, are specified. (B) Biological processes enriched in modules distinct to G3 and G4/rhabdoid cells with at least 10 members following decomposition of the PPI networks in (A) based on KEGG and Reactome annotations. Enriched terms were determined by an FDR < 0.05 (Fisher’s exact test with Benjamini-Hochberg correction). Only non-redundant terms are presented, except where additional terms provided complementary information. The fraction of proteins in each module with the corresponding annotation is provided.

Pathways associated with upregulated elements provided important and novel insights into the biology of rhabdoid differentiation. For instance, analysis of cellular regulators that were identified with a high centrality in the network constructed for cells with rhabdoid differentiation highlighted Forkhead box transcription factor 1 (FOXM1) as a prominent hub in Module IV (Fig. 5A). As FOXM1 plays a critical role in regulating the cell cycle, it is implicated in the control of genes essential for cell proliferation and progression (TRRUST v2)^36^. Of note, this node was of much lower significance in low- and high-grade ccRCC networks (Fig. 5A and Suppl. Fig. 4A) indicating an elevated potential for proliferation in cells with the rhabdoid phenotype. Importantly, high FOXM1 expression was associated with poor prognosis in the TCGA-KIRC cohort (Suppl. Fig. 4B) underscoring the relevance of its overexpression for patient prognosis. Additionally, the cell cycle subnetwork in rhabdoid cells encompassed several molecules that function downstream of the FOXM1 transcription factor such as DNA replication factors (CLSPN, CDC6), elements implicated in the G1/S- (CDC25A, CCNA2, CDK3) and G2/M-checkpoints (PLK1, CDC25A/C, BUB1) (Fig. 5A, 5B and Suppl. Fig. 4C). Their dysregulation likely contributes to the increased proliferative potential of rhabdoid ccRCC cells. Additionally, in this subnetwork, we observed rhabdoid ccRCC cell-specific upregulation of BIRC5 (Fig. 5B), a well-established inhibitor of apoptosis protein (IAP).

**Figure 5:**
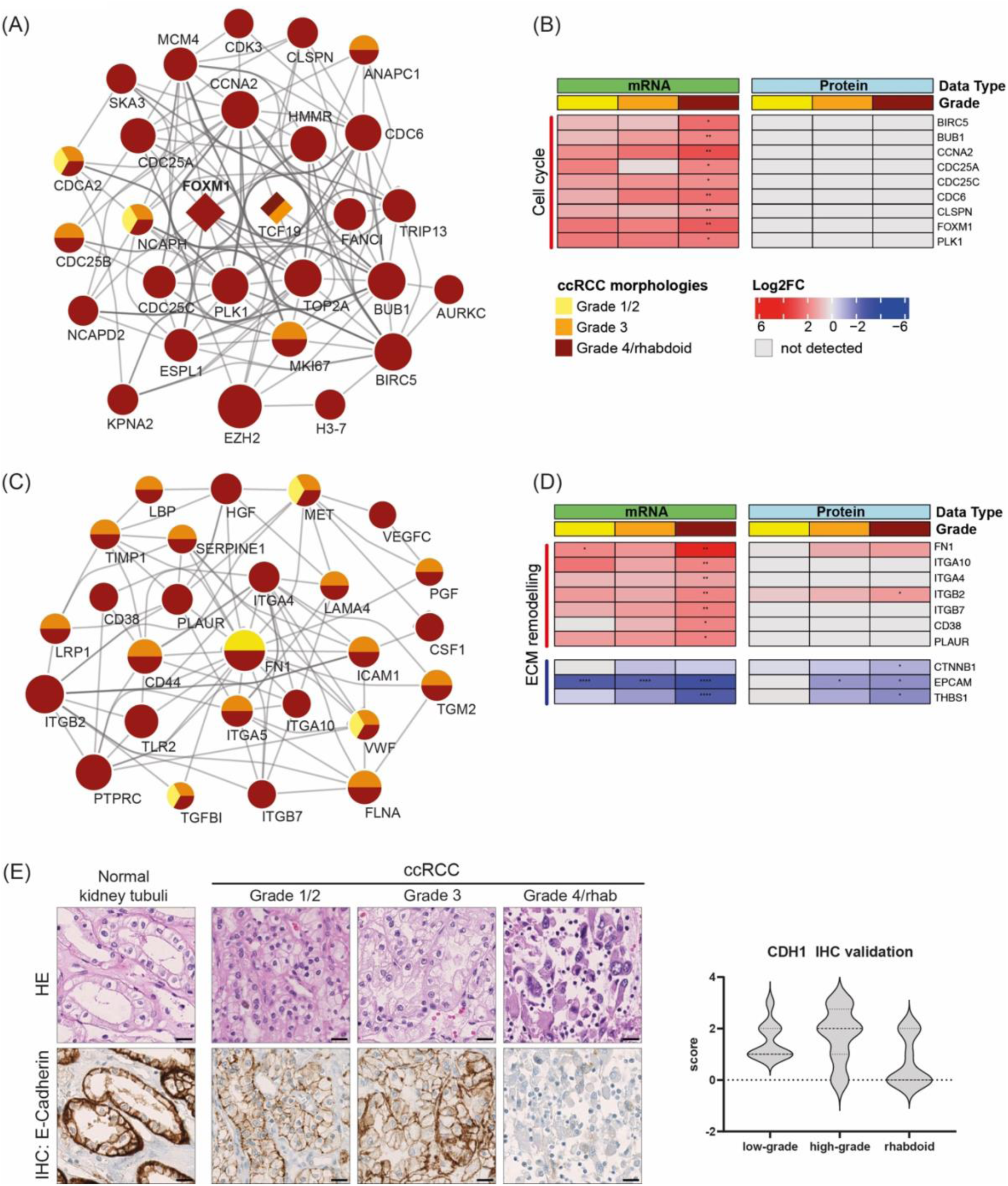
Grade 4/rhabdoid ccRCC cells are characterized by enhanced expression of cell cycle factors and dysregulation of cell-matrix interactions. (A) Network decomposition highlighting a subnetwork associated with upregulation of cell cycle factors (Reactome R-HSA-1640170) in the grade 4/rhabdoid ccRCC cells (Module IV). Factors that are significantly upregulated also in lower grade ccRCC cells are indicated. Bigger nodes correspond to higher centrality values in the Grade4/rhabdoid ccRCC PPI network. Grades are color-coded as in (B). (B) Heat maps highlighting selected elements from module IV and their respective dysregulation during ccRCC de-differentiation on both data layers. Alterations of representative genes and proteins are depicted as log2FC with respect to adjacent normal cells. FDR is indicated by “****” <= 0.0001, “***” <= 0.001, “**”<= 0.01, “*” <= 0.05. (C) Network decomposition highlighting a subnetwork containing upregulated elements involved in focal adhesion (KEGG path:hsa04510), integrin cell surface interactions (Reactome R-HSA-216083), homeostasis (Reactome R-HSA-109582) and proteoglycans implicated in cancer (KEGG path:hsa05205) in the grade 4/rhabdoid ccRCC cells (Module V). Factors that are significantly upregulated in lower grade ccRCC cells are indicated. Bigger nodes correspond to centrality values in the Grade4/rhabdoid ccRCC PPI network. Color codes as in (B). (D) Heat maps highlighting selected elements implicated in ECM remodelling and cell adhesion and their respective dysregulation during ccRCC de-differentiation on both data layers. Alterations of representative genes and proteins are depicted as log2FC with respect to adjacent normal cells. Color codes are depicted in (B). FDR is indicated by"****" <= 0.0001, "***" <= 0.001, "**"<= 0.01, "*" <= 0.05. (E) IHC analysis of E-cadherin expression ccRCC with distinct grade morphologies. Representative images are shown for each grade group, the scale bar represents 20 mm. On the right: Expression levels were scored from 0 (no expression) to 3 (high expression) and the quantification of N=11 cases is shown.

Module V in the G4/rhabdoid-specific network encompassed elements involved in focal adhesion, integrin cell surface interactions, homeostasis and proteoglycans implicated in cancer (Fig. 5C, Suppl. Table 10) thus emphasizing the significance of tumor-cell driven re-organization of the extracellular matrix (ECM) for ccRCC progression and metastatic dissemination. This module contained several genes that were specific for the grade 4/rhabdoid network and that had high centrality scores. Among these, were integrins (ITGA4, ITGA10, ITGB2, ITGB7) with known roles in influencing tumor growth, metastasis, and the interactions within the tumor microenvironment (TME)^37^. Importantly, among these factors ITGA4 was associated with poor prognosis in the TCGA-KIRC cohort (Suppl. Fig. 5A). Further nodes in this module included other upregulated elements with known functions in modulating the TME, such as FN1, PLAUR, VEGFC and CD38. These factors have been linked to EMT thereby promoting cancer cell migration and invasion. Conversely, we observed downregulation of THBS1, CDH1 and EPCAM specifically in the rhabdoid network (Fig. 5D and Suppl. Table 10) underscoring the overall dysregulation of cell-matrix interactions and enhanced activation of EMT. Immunohistochemistry (IHC) analysis confirmed an almost complete absence of E-cadherin protein expression in the rhabdoid component of morphologically heterogeneous ccRCC cases, suggesting its potential utility as a companion biomarker to aid accurate diagnosis of grade 4 rhabdoid differentiation (Suppl. Table 8 and 9, Fig. 5E). The relevance of this finding was further underscored by the association of low CDH1-mRNA expression with poor survival in the TCGA KIRC cohort (Suppl. Fig. 5B).

These results not only shed light on the specific mechanisms contributing to the malignant potential of rhabdoid ccRCC cells, but they also highlight the molecular alterations underlying their structural hallmarks, such as large cells with discohesive growth patterns and high mitotic activity, which are likely driven by altered cell adhesion and enhanced proliferation.

### Immune evasion and dysregulation are hallmarks of rhabdoid ccRCC cells

Our analysis of transcriptome and proteome data suggested that immune response dysregulation was specific to rhabdoid ccRCC cells across both data layers (Fig. 2E and 2F, Fig. 3B). To investigate this further, we analyzed CD8+ T-cells infiltration in a larger cohort of ccRCCs with grade 4/rhabdoid areas. We detected CD8+ T-cells in almost all grade 4/rhabdoid ccRCC cases of our extended cohort (Suppl. Table 8 and 9) indicating inflamed phenotypes of varying degrees (Fig. 6A). In our grade4/rhabdoid-specific mRNA expression data we also observed significant upregulation of the immune checkpoint molecule PD-L1 (CD274). Of note, this was not the case for other well-studied checkpoint factors such as CTL-4 or LAG3 (Suppl. Fig. 5C). Integrated network analysis highlighted an immunosuppressive network largely specific to grade 4/rhabdoid ccRCC cells, which was distinct from lower-grade cells (Modules III, XI and XXIII) (Fig. 6B). In these modules, we found high expression of chemokines including CCL4, CCL20, CXCL1 and CXCL13 (Fig. 6C), molecules that can interact with the TME and immune cells to foster an immunosuppressive milieu^38–40^. In addition, elevated type I interferon (IFN) signaling, particularly IFN-alpha and IFN-beta, emerged as a prominent feature in module III. High expressions of STAT1/2, the main signaling molecules of the IFN pathway, was associated with grade 3 and grade 4/rhabdoid cells (Fig. 6B and 6C). Upregulation of STAT1/2 transcriptional targets were predominantly enriched for interferon-beta signaling molecules (OAS1, MX1 and IFI44) (Fig. 6B and 6C, Suppl. Fig. 4C)^41^. Highly interestingly, we also identified upregulation of CD38 in grade 4/rhabdoid ccRCC cells, a known target of IFN-beta signaling (Fig. 5D). CD38 associated with the cell adhesion network module in our integrated network analysis (Module V, Fig. 5C) but was more recently described to suppress cytotoxic T cells activity when overexpressed on tumor cells^42^ suggesting CD38 overexpression may have a potential role in immune evasion in rhabdoid ccRCC. Additionally, we observed CD38 upregulation in RNAseq data from cases with rhabdoid differentiation in the CheckMate 025 cohort, irrespective of the treatment arm (Fig. 6D). However, due to the limited number of samples with rhabdoid differentiation and accompanying RNAseq data in the nivolumab arm (N = 8), it was not feasible to draw definitive conclusions regarding the clinical benefit of immune checkpoint inhibitors (ICI) in this subgroup. Besides CD38, also ITGB2, another factor with roles at the intersection of ECM remodeling and immune modulation, showed prominent upregulation in grade 4/rhabdoid ccRCC cells (Fig. 5C and 5D) likely contributing to a state of immune suppression in rhabdoid ccRCC^43^.

**Figure 6:**
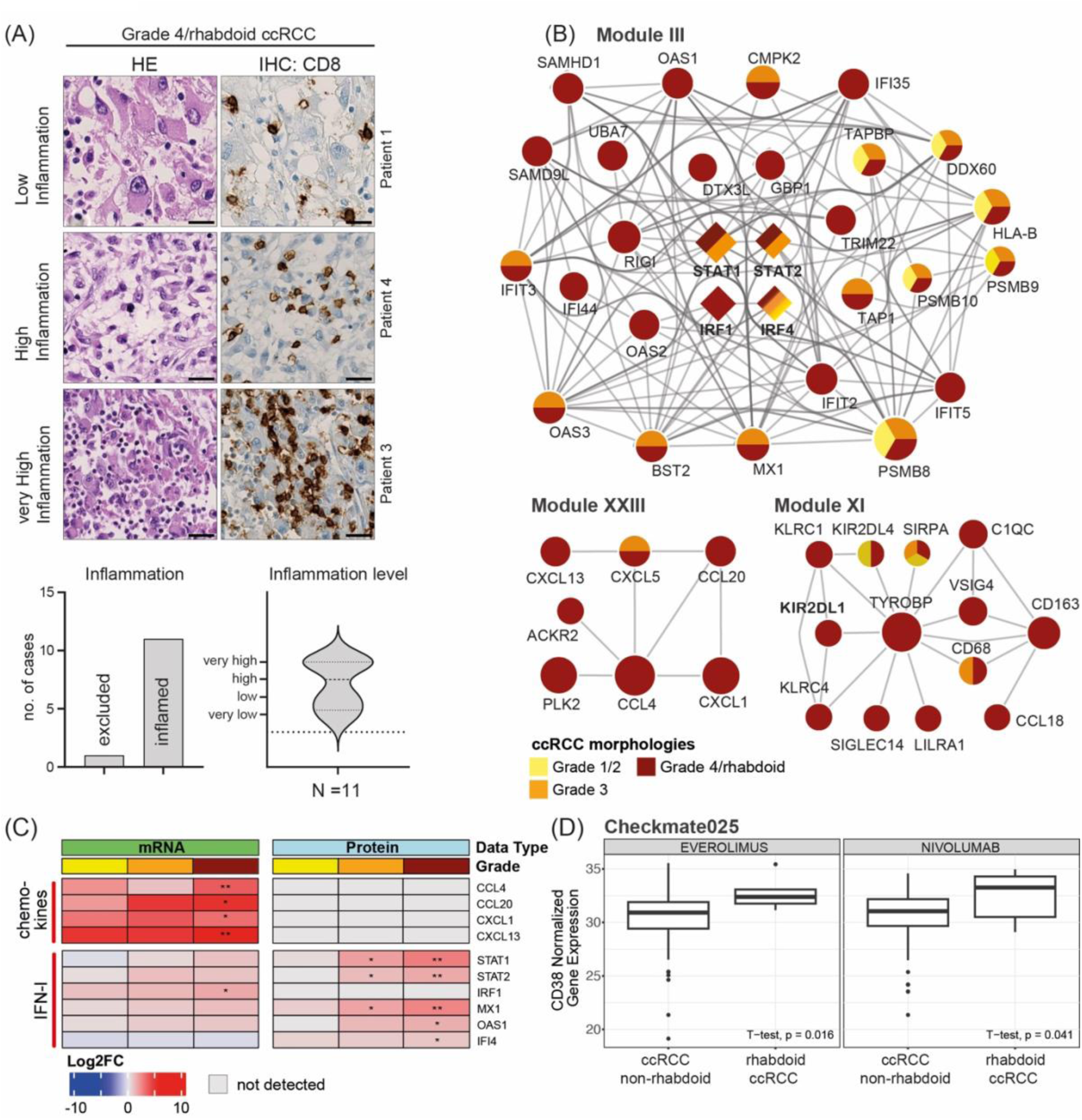
Aggressive ccRCC cells show upregulation of the immune response. (A) IHC analysis using antibodies against CD8 depicting infiltration of CD8+ T cells into the rhabodoid component of the ccRCC cases in this study. CD8+T cells were scored and assigned to the depicted immune phenotypes (inflamed or excluded) by pathologist review. Inflamed tumors were further categorized into low, high or very high inflammation levels and representative images for each phenotype are shown. The scale bar represents 20 mm. We analyzed 12 cases with rhabdoid de-differentiation and quantified the frequency of inflammation (CD8+ T cells) as well as the level (low, high, very high) focussing on the rhabdoid component of the ccRCCs. (B) Integrated network analysis showed a significant enrichment of upregulation of immune response elements in Modules III (Interferone Signaling, Rectome R-HSA-913531 and Immune System, Reactome R-HSA-168256), XI (Immune System, Reactome R-HSA-168256) and XXIII (Chemokine receptors bind chemokines, R-HSA-380108) of the grade4/rhabdoid ccRCC network. Factors that are significantly upregulated also in lower grade ccRCC cells are indicated. Bigger nodes correspond to higher centrality values in the grade4/rhabdoid ccRCC PPI network. (C) Heat maps highlighting selected elements associated with the immune system-related pathways and their respective dysregulation during ccRCC de-differentiation on both data layers. Alterations of representative genes and proteins are depicted as log2FC with respect to adjacent normal cells. FDR is indicated by"****" <= 0.0001, "***" <= 0.001, "**"<= 0.01, "*" <= 0.05. (D) Boxplots of normalized CD38 gene expression in non-rhabdoid vs. rhabdoid ccRCC patients from the CM-025 clinical trial cohort for the everolimus (Non-rhabdoid ccRCC N=108, Rhabdoid ccRCC N= 6) as well as nivolumab (Non-rhabdoid ccRCC N=139, Rhabdoid ccRCC N= 8) arms. Significance was tested using a Welch t-test.

Take together, our analysis revealed a substantial immunosuppressive network driven by signaling pathways activated specifically in rhabdoid ccRCC cells. Upregulated factors like CD38 and ITGB2 appear to exert immunomodulatory functions that inhibit the activity of abundant CD8+ T-cells within the rhabdoid ccRCC TME.

## Discussion

Morphological intra-tumor heterogeneity (ITH) is a well-documented phenomenon, where for example different grade morphologies can co-occur within the same tumor section^44,45^. Modern microscopy combined with AI-based analysis methods has been successfully applied in biomedical imaging for segmentation and classification tasks across various cancer types^46–50^. Our AI-driven automated classification system reliably grades and classifies ccRCC cells in HE-stained digital WSI and thus enhances the accuracy of histopathological assessments and helps to overcome the need for manual expert annotations.

Previous studies have shown the profound genetic ITH in ccRCC tissue samples^5,51^ but a correlation with morphological features was not possible with their approaches. Combining AI-based image analysis with single-cell isolation and deep multi-omics profiling of selected cells provided us with unprecedented insights into the relationship between molecular and morphological heterogeneity in ccRCC. This revealed a trajectory of progressive dysregulation in ccRCC cells with increasing grades on both, mRNA and protein levels, indicating that non-genetic alterations likely contribute to their malignancy. Importantly, mRNA and protein measurements provided complementary information and consistent with previous data, we observed that overall downregulation of pathways was more prominent than upregulation during ccRCC tumor cell differentiation^32,52^. As tumor grade increased, ccRCC mRNA and protein data showed overlapping functional alterations, highlighting progressive dysregulation of metabolic pathways such as one-carbon metabolism and decreased expression of proximal renal tubule genes essential for kidney function^32^. Notably, grade 4/rhabdoid ccRCC cells appeared to diverge most from highly differentiated kidney tubular cells, showing increased expression of stem cell markers like CD44 and DCLK1^33,34^, implying a potential role of stem cell-like features for the aggressiveness in high-grade ccRCC. As the acquired expression profiles reflected many known ccRCC tumorigenic changes^6,52^, we were confident in the accuracy of mRNA and protein dysregulation portraits obtained from as little as 1000 individually collected ccRCC cells from FFPE sections.

While recent studies have explored the molecular landscape and therapeutic vulnerabilities of RCC with sarcomatoid differentiation^13,16^, the biology of rhabdoid ccRCC remained understudied^14^. Bakouny and colleagues profiled the molecular features in grade 4 ccRCC cells but did not identify molecular characteristics that distinguished rhabdoid differentiation in RCCs^13^. However, the majority of cases of their cohort comprised sarcomatoid RCCs, raising the question whether rhabdoid ccRCC is truly similar to or biologically distinct from sarcomatoid cases, as their morphologies suggest. Previous molecular investigations have associated the rhabdoid component with BAP1 mutations^13,45^. However, our genome-wide genetic profiling of five ccRCC tissue samples with rhabdoid differentiation did not corroborate that BAP1 mutations were a feature in most cases. We also did not observe the loss of chromosome 11p, which was previously reported as a distinct feature in ccRCC with rhabdoid differentiation^53^. Instead, our study highlights specific transcriptome and proteome alterations in a homogeneous cohort of ccRCCs with rhabdoid differentiation. We identified upregulation of FOXM1-driven proliferative signaling and ECM remodeling as key molecular changes associated with high-grade and rhabdoid phenotypes. Interestingly, FOXM1 has been associated with tumor progression and poor prognosis in patients with ccRCC before^54^. Upregulations of integrins on the other hand, was shown to enhance tumor cell motility and interactions within the ECM, thus promoting tumor growth and metastatic dissemination, potentially by modulating matrix metalloproteinases (MMPs)^37^. Of note, integrins have been associated with worse prognosis in ccRCC and particularly ITGB2 was shown to be overexpressed in aggressive cancers^55–57^. On the other hand, enhanced proliferation and ECM remodeling likely explain the distinct morphological features observed in rhabdoid cells in ccRCC as these processes disrupt normal cell adhesion and structural integrity within the tumor, allowing cells to assume a more invasive, migratory phenotype^20^. Here we highlight distinct molecular alterations in rhabdoid ccRCC cells and suggest that targeting these networks could be crucial in preventing metastasis and improving patient survival.

Moreover, our study unveils a distinct immunosuppressive signaling network driven by rhabdoid ccRCC cells, involving well-known immune checkpoints like PD-L1 and novel immunomodulatory factors such as CD38 and ITGB2^43,58,59^. Interestingly, ITGB2, which we found overexpressed in rhabdoid ccRCC cells, has recently been identified to regulate the immune response within the TME besides its more established roles in the integrin-ECM signaling network that promote EMT and metastatic spread^43,60^. Additionally, elevated interferon signaling, chemokine, and integrin expression further characterized this immunosuppressive network. Consistently, we observed high STAT1/2 expression and elevated IFN-I signaling in rhabdoid ccRCC cells which likely drive EMT, immune evasion, and an aggressive phenotype in inflammatory tumors^41,61^. STAT1/2 in particular are well recognized markers of aggressiveness in ccRCC^32^. These findings suggest that rhabdoid ccRCC cells play key roles in reshaping the TME, creating an immunosuppressive milieu that likely compromises CD8+ T-cell function and may mediate resistance to ICI^58^. These findings were further corroborated by two recent studies that highlight the significant role of the TME for ccRCC progression and advances disease showing that these exhibit enrichment of terminally exhausted CD8+ T cells^62,63^

Although clinical inter-patient heterogeneity and the limited number of samples pose challenges, our study is the first to integrate AI-driven image analysis, single-cell isolation, and deep multi-omics profiling to characterize the molecular aberrations in distinct clinically relevant and morphologically defined aggressive cell populations. Analysis of larger cohorts, such as the CheckMate 025 trial^64^, revealed comparably low numbers of rhabdoid ccRCC samples with accompanying molecular data. Despite this limitation, our analysis of RNAseq data from the trial demonstrated elevated CD38 expression in rhabdoid ccRCCs, highlighting its potential relevance. However, the small sample size precluded any conclusions regarding the clinical benefit of ICIs in this subgroup. Taken together, our findings offer valuable insights into the molecular pathways underpinning grade 4/rhabdoid ccRCC, including FOXM1-driven cell cycle regulation, ECM remodeling, and immune evasion. These data provide a strong rationale for exploring combination therapies targeting these pathways to enhance the efficacy of ICIs^59^ in this challenging patient population.

## Methods

### RESOURCE AVAILABILITY

#### Lead contact

Further information and requests for resources and reagents should be directed to and will be fulfilled by the lead contacts, Dr. Hella A. Bolck and Prof. Dr. Peter Horvath.

#### Materials availability

This study did not generate new unique reagents.

#### Data and code availability

##### Data Availability

Patient and clinical data related to analyses in this study are listed in Supplementary Tables 1, 4 and 8. Histological images used for automated image analysis and cell classification are deposited in FigShare using the following link: https://doi.org/10.6084/m9.figshare.c.7682516. A demo dataset and online material of how to install BIAS and reproduce our work can be accessed at https://drive.google.com/drive/folders/13QlATL8r9TW47tDxjnJ2owfNT4C2PwOs?usp=drive_link.

Additional details are provided in Supplementary Table 4. Whole genome sequencing (WGS) and RNAseq data are not publicly available due to restrictions concerning privacy of research participants and can be made available from the corresponding authors upon reasonable request. Filtered somatic mutation data are available in Supplementary Tables 2 and 3. RNAseq read count data have been deposited in Zenodo together with the accompanying code https://doi.org/10.5281/zenodo.14922388 (please find the fully tokenized URL in the section “Code”). The mass spectrometry proteomics data have been deposited to the ProteomeXchange Consortium via the PRIDE^65^ partner repository with the dataset identifier PXD061011. Details regarding data collection are included in Supplementary Table 5. Results from overrepresentation analysis are available in Supplementary Tables 6 and 7. Immunohistochemistry scores are detailed in Supplementary Table 9 with corresponding raw images available upon reasonable request from the authors. Data from the CheckMate 025 clinical trial is available as part of the accompanying publication^64^.

##### Code availability

A free compiled version of BIAS with limited high-throughput capabilities is available at https://drive.google.com/drive/folders/13QlATL8r9TW47tDxjnJ2owfNT4C2PwOs?usp=drive_link. containing all features applied in the described workflows. Algorithms used for omics data analysis are publicly available from the indicated references in the Methods section. Specialized code used for RNAseq and proteomics data analysis in this study were deposited in Zenodo: https://zenodo.org/records/14922388?token=eyJhbGciOiJIUzUxMiJ9.eyJpZCI6IjUzYWMyZTBmLTJkNmUtNGFmNy1hZTY0LTVlZWI1ODI2NTdmZCIsImRhdGEiOnt9LCJyYW5kb20iOiJhNTdmZTg5MTQzMTliNmQ2MThiNzkwNmViYTM0ZTlhZSJ9.ZX1CaI1ri1IwsbeJvWqSxVZqDnO_B9fO94HOUcOadfKX-o2aBZUsdYfQCIQpfDMG1b9Ord97f3L6hlB8UL2WPQ (full tokenized URL).

The code for multi-omics data integration analysis as well as for automated network representations in Cytoscape can be found on GitHub at https://github.com/TotuTiberiu/NOODAI and corresponds to the version 1.0.0 of the NOODAI platform^35^ that can be freely accessed at https://omics-oracle.com/NOODAI including instructions and tutorials).

### EXPERIMENTAL MODEL AND STUDY PARTICIPANT DETAILS

#### Patient cohort

The Department of Pathology and Molecular Pathology at University Hospital Zürich made available archival FFPE tissue samples for this study. It was approved by the local Ethics Committee (BASEC# 2019-01959) and in agreement with Swiss law (Swiss Human Research Act). All patients gave written consent. The tumor samples used in this study were diagnosed as ccRCC and the overall grade was noted from the diagnostic histopathologic patient reports according to the WHO/ISUP^9^. Two board-certified pathologists re-assessed all FFPE tissue samples in the context of this study to confirm the presence of malignant and normal areas as well as different grade morphologies. Associated clinical information is shown in Suppl. Table 1, 4 and 8.

### METHOD DETAILS, QUANTIFICATION AND STATISTICAL ANALYSIS

#### Genomic characterization by whole-genome sequencing (WGS)

To extract tumor and normal DNA from our core cohort, we sampled all areas that were included in the image analysis and single-cell isolation workflow to cover all mutations that may be present at these sites. To facilitate this a 2 µm section was stained with hematoxylin and eosin (H&E) to evaluate the morphology and an experienced pathologist marked the areas for dissection (Suppl Fig. 1). This slide was used as a reference image for the AVENIO Millisect System (Roche) and subsequently the marked areas were dissected from a consecutive unstained 10 µm section prepared on a glass slide. Tissue areas from at least two 10 µm sections were collected in a tube containing lysis buffer (Promega, A826E from Maxwell® 16 FFPE Tissue LEV DNA Purification Kit) and subjected to genomic DNA extraction on a Maxwell® 16 machine (Promega) following the manufacturer’s instructions.

Whole-genome sequencing libraries were constructed according to Illumina TruSeq DNA PCR Free Library Prep protocol HT (Illumina Inc., San Diego, CA, USA). This included: (1) fragmentation of 400 ng genomic DNA to 350 bp inserts by Covaris LE220-plus, (2) cleanup of fragmented DNA, (3) end repair, (4) removal of large and small DNA fragments, (5) 3′-end adenylation and (6) adapter ligation. The resulting libraries were quantified and quality-assessed with the iSeq100 (Illumina). The samples were sequenced with the NovaSeq 6000 platform (Illumina) using S4 flow cells with 300 cycles (2 × 150 reads) and measured 2 × to reach an average coverage of 30X for normal kidney tissue and 120X for the tumor fraction per patient.

The raw sequencing files (bcf files) were converted to fastq format and demultiplexed in a single step using Illumina’s bcl2fastq program via Illumina DRAGEN Bio-IT Platform (v3.8.4). First steps in secondary analysis, including read trimming, alignment to GRCh38, sorting, duplicate marking and somatic variant calling were done using Illumina DRAGEN Bio-IT Platform (version 3.9.5) ^66^ or Mutect2 ^67^ and Strelka2 ^68^. To identify and annotate somatic variants, we annotated and converted the VCF files into MAF format using VEP ^69^. The MAF files were analyzed using Maftools in R ^70^. In addition, Maftools was used for comprehensive analysis and visualization, including tumor mutational burden (TMB) assessment, mutational signature analysis, and identification of significantly mutated genes. Genome-wide copy number aberrations were visualized with gistic (v2.0.23) using default parameters and log-transformed segmentation files from DRAGEN as input.

#### Sample preparation for morphological analysis

For each ccRCC case from our core cohort, we selected one to four representative FFPE blocks in order to include areas displaying low-grade, high-grade and rhabdoid areas as well as normal kidney histology. Subsequently, 10 µm sections were prepared using PEN MembraneSlides 1.0 (Carl Zeiss) and stained with H&E. Slides were digitized scanning z-stacks with seven z-slices in 0.5 µm increments using a Hamamatsu C9600 scanner at ×40 optical magnification. The scanner resolution was 0.228 µm per pixel. The NDPI files generated by the Hamamatsu C9600 scanner were processed into a BIAS-compatible format as follows. First, the NDPI split tool was used to extract image data and generate non-overlapping tiles with a resolution of 1024 × 1024 pixels. Tiles were saved in TIFF format, preserving spatial information. After tile extraction, the Simple Focus tool was applied to automatically select the optimal focus plane. The algorithm computes a sharpness value for each image field with seven Z positions based on four-neighborhood pixel differences. These differences are accumulated into a single sharpness score, and the field with the highest sharpness value was selected.

#### Image analysis

Using the BIAS software suite^30^, we developed a pipeline to assign every cell to one of the following classes (i) grade 1/2 (low-grade) ccRCC, (ii) grade 3 (high-grade) ccRCC, (iii) grade 4/rhabdoid ccRCC, (iv) normal kidney, (v) other/junk. This workflow was composed of several steps: segmentation of nuclei, specification of cytoplasmatic areas, pre-classification into tumor and normal kidney cells, segmentation of nucleoli, segmentation of rhabdoid cells, classification into the distinct classes (i-v) mentioned above. We used Artificial Neural Networks (ANN) for both segmentation and classification. The classification models were multilayer perceptron networks (MLPs) trained on features extracted from the segmented cells. To segment nuclei, we integrated nucleAIzer models into BIAS^31^. Dilating the nuclear contours by 10 µm created the cytoplasmatic masks that were used to extract features from the cytoplasm. These were aggregated with features of the 10 neighboring cells and an artificial neural network pre-classified the tissue into healthy and tumor.

Instance segmentation algorithms based on a Mask-RCNN implementation with a Resnet101 backbone achieved custom segmentation of prominent nucleoli and rhabdoid cells. Additional cases were annotated as training data (Suppl. Table 4). Subsequently, cRCC cells were classified into grade 1/2, grade 3, grade4/rhabdoid, normal kidney or other/junk by MLP classification models trained on features extracted from the segmented cells. Phenotype predictions were ranked by confidence and the 2000 best cells were manually curated. These were used as an extended training set to improve the ML classification further.

K-fold stratified cross-validation (where K=10) was performed using the core patient cohort (Suppl. Table 1). Moreover, we evaluated our approach using six independent grade 4 ccRCC cases with rhabdoid morphology and grade ITH (Supp. Table 8).

#### Sample preparation and single cell isolation

H&E-stained FFPE tissue sections on PEN MembraneSlides were subjected to coverslip removal by immersing them in xylene for 1-2 hours (until the coverslip loosened) followed by xylene removal and hydration using descending ethanol concentrations and air-drying. Subsequently, cells were isolated by LMD based on the morphologies assigned by the machine learning classification module of BIAS in the same sections. Relevant cell contours were transferred to a Leica LMD6 microscope for automated single-cell isolation. The Leica LMD6 system, equipped with an HC PL FLUOTAR L ×63/0.70 objective, was used for cell excision. Cells were excised based on nuclei contours plus a 6 µm offset. For each morphological class in each sample, two sets of 1,000 cells were microdissected and collected into PCR caps (VWR, 732-0548) or 384-well plates (Eppendorf® Twin tec PCR, 384 wells EP0030128508) To cells isolated for RNAseq analysis, 5 μl of PKD buffer (Qiagen) and 25 μl of RNA later (Qiagen) were added, followed by centrifugation at 10,000 g for 5 minutes. For proteome analysis we added 10μl of ammonium bicarbonate buffer to the isolated cells.

#### RNA sequencing and data analysis

RNA was extracted from microdissected single cells using the RNeasy FFPE kit (Quagen, Hilden, Germany). Entire RNA extracts were transferred into NGS Library preparation without QC. Due to the low concentration of total RNA the SMARTer® Stranded Total RNA-Seq Kit v2 -Pico Input Mammalian (A Takara Bio Company, California, USA) was used to prepare sequencing libraries.

Briefly, total RNA samples were reverse-transcribed using random priming into double-stranded cDNA in the presence of a template switch oligo (TSO). When the reverse transcriptase reached the 5’ end of the RNA fragment, the enzyme’s terminal transferase activity added non-templated nucleotides to the 3’ end of the cDNA. The TSO paired with the added non-templated nucleotide, enabling the reverse transcriptase to continue replicating to the end of the oligonucleotide. This resulted in a cDNA fragment that contained sequences derived from the random priming oligo and the TSO. PCR amplification using primers binding to these sequences could then be performed. The PCR added full-length Illumina adapters, including dual barcodes for multiplexing. Ribosomal cDNA was cleaved by ZapR in the presence of the mammalian-specific R-Probes. Remaining fragments were enriched with a second round of PCR amplification using primers designed to match Illumina adapters (16 cycles). The quality and quantity of the enriched libraries were validated using the Tapestation (Agilent, Waldbronn, Germany). The product was a smear with an average fragment size of approximately 300 - 360 bp. The libraries were normalized to 5nM in Tris-Cl 10 mM, pH8.5 with 0.1% Tween 20.

A shallow sequencing approach on the iSeq (Illumina, Inc, California, USA) was used to check the read distribution among samples, rRNA content and reference genome mapping. The Novaseq 6000 (Illumina, Inc, California, USA) was used for deep sequencing according to the standard manufacturer’s protocol. Sequencing configuration was single-end 100 bp. Demultiplexing was performed with bcl2fastq (Illumina, Inc, California, USA) allowing for 0 mismatch on Index 1 and 1 mismatch on Index 2. Between 70 million and 146 million reads were generated per sample.

RNA sequencing data analysis was performed using the SUSHI framework^71^, which encompassed the following steps: Read quality was inspected using FastQC, and sequencing adaptors removed using fastp; Alignment of the RNASeq reads using the STAR aligner^72^ and with the GENCODE human genome build GRCh38 (Release 42) as the reference^73^; the counting of gene-level expression values using the ‘featureCounts’ function of the R package Rsubread^74^; differential expression using the generalized linear model as implemented by the DESeq2 Bioconductor R package^75^, and; Gene Ontology (GO) term pathway analysis using the hypergeometric over-representation and GSEA tests *via* the ‘enrichGO’ and ‘gseGÒ functions respectively of the clusterProfiler Bioconductor R package^76^. For differential gene expression analysis (DEG) we contrasted gene expression in each ccRCC grade to that in the normal renal tubuli samples to obtain three ccRCC-specific comparisons termed (i) grade 1/2 (low-grade), (ii) grade 3 (intermediate/high-grade, (iii) grade 4/rhabdoid. Additional visualizations were generated in the exploreDE Interactive Shiny app^77^. All R functions were executed on R version 4.2.0 (R Core Team, 2020) and Bioconductor version 3.15.

#### Proteomics data acquisition and analysis

For sample lysis, the collected cells were spun down and 20 μl of 50 mM Triethylammonium bicarbonate (TEAB) were added to each tube or well. The 384-well plate was closed with sealing tape (Corning, CLS6569-100EA) and centrifuged at 2000 g for 2 min. All samples were incubated for 1 hour at 95 °C. For the plates a PCR thermo cycler with 384-well reaction module (Biorad S1000) was used, for the PCR tubes, the 96-well module (Eppendorf, Mastercycle X50a) was installed. The lid temperature for all incubation step was always 110 °C. Prior to incubation for 60 min at 75°C, 5 µl of 60 % acetonitrile in 50mM TEAB was added. For in-solution protein digestion, samples were cooled to room temperature and subjected to pre-digestion using 1 µl LysC (4 ng/µl) for 4 h at 37 °C. Subsequently, 1 µl Trypsin (4 ng/µl in TEAB) was added for overnight digestion at 37 °C. In the morning, another 4 ng of Trypsin was added and the samples were incubated for 5 hours at 37 °C^78^. The digestion was stopped by acidification, adding 10 µl 5 % aqueous trifluoroacetic acid and the sample were stored at −20 °C until LC-MS analysis.

For proteomics data acquisition, **h**alf of the digested samples were loaded on Evotips (Evosep Biosystems) following the provided instructions and analysed on an Evosep One LC coupled to a TIMS TOF Pro mass spectrometer (Bruker). The mass spectrometry proteomics data were handled using the local laboratory information management system b-fabric (LIMS) ^79^, and all relevant data have been deposited to the ProteomeXchange Consortium via the PRIDE (http://www.ebi.ac.uk/pride) partner repository with the data set identifier [tbd].

The acquired data-independent-acquisition (DIA) spectra were processed with DIA-NN (Version 1.8.1.8)^80^, using a library free approach with the ensembl homo_sapiens data base (release-108) and oxidation at methionine as variable modification. The R package prolfqua^81^ was used to process, handle and analyze the differential expression and to determine group differences, confidence intervals, and false discovery rates for all quantifiable proteins. Starting with the report.tsv files generated by DIA-NN, which included the precursor ion abundances for each raw file, we determined gene centric abundances by first aggregating the precursor abundances to peptidoform abundances after filtering for gValues < 0.01. Then, we employed the Tukeys-Median Polish to estimate gene abundances. This was possible with a mapping table where for each ensemble protein the gene name was listed (if there was no gene name listed, the ensemble protein identifier was used). Using the gene names instead of the protein identifier allowed us to roll-up the peptidoform abundances to gene names. Furthermore, before fitting the linear models, we transformed the gene abundances using the log2 transformation followed by robust scaling normalization.

For differential protein expression analysis (DEP) we contrasted protein expression in each ccRCC grade to that in the normal renal tubuli samples to obtain three ccRCC-specific comparisons termed (i) grade 1/2 (low-grade), (ii) grade 3 (intermediate/high-grade, (iii) grade 4/rhabdoid. Gene Ontology (GO) term pathway analysis was performed in the same way as in the RNA sequencing data analysis pipeline using the hypergeometric over-representation and GSEA tests *via* the ‘enrichGO’ and ‘gseGÒ functions respectively from the clusterProfiler Bioconductor R package^76^.

#### Immunohistochemistry (IHC)

Sections (2 µm) were prepared from FFPE specimens and were stained with haematoxylin and eosin (H&E) for histological evaluation. For IHC, sections (2 μm) were transferred to glass slides followed by antigen retrieval. IHC was performed using the Ventana Benchmark automated system (Roche, Ventana Medical Systems, Oro Valley, AZ) and Ventana reagents. The Optiview DAB IHC detection kit (Roche, Ventana Medical Systems) was used to stain with the antibody against CD8 (Clone C8/144B, cat. number M7103, DAKO, used 1:100). The staining procedure for CDH1 (E-cadherin clone 36, cat. number 610181, BD Biosciences, used 1:500) was carried out with the automated Leica BOND system using the Bond Polymer Refine Detection Kit (Leica Biosystems, Wetzlar, Germany).

#### Survival analysis

Survival curves were generated using the Gene Expression Profiling Interactive Analysis (GEPIA) web tool (http://gepia.cancer-pku.cn) to assess the prognostic significance of gene expression levels for CDH1, FOXM1 and ITGA4 in the TCGA Kidney Renal Clear Cell Carcinoma (TCGA-KIRC) patients. Patients were divided into high and low expression groups based on the median expression level. Kaplan-Meier survival curves were generated, and the log-rank test was used for statistical comparison. Hazard ratios (HR), 95% confidence intervals (CI), and p-values were calculated and displayed. Survival differences were considered significant if p-values were <0.05. HRs >1 indicated higher risk, while HRs <1 indicated a protective effect.

#### Statistical testing

We performed differential expression analysis (DEG/DEP) comparing ccRCC grade groups (grade 1/2, grade 3, grade 4/rhabdoid) to adjacent normal kidney tubule cells. To assess significant changes, we applied a false discovery rate (FDR) threshold of < 0.05 and considered log2 fold change (log2FC) greater than |1|. The significance levels for FDR are denoted as follows: **** <= 0.0001, *** <= 0.001, **<= 0.01, * <= 0.05. Additionally, we color-coded the log2FC values for visual representation.

For enrichment analysis in the transcriptomic and proteomic datasets, we conducted over-representation analysis using the hypergeometric test. Specifically, we examined Gene Ontology (GO) biological pathways. Benjamini-Hochberg adjusted p-values were calculated and Significance levels are denoted as follows: **** for p <= 0.0001, *** for p <= 0.001, ** for p <= 0.01, * for p <= 0.05.

Patient and upper-quartile (UQ) normalized transcripts-per-million (TPM) RNAseq data from CheckMate 025 (CM-025; NCT01668784) prospective clinical trial of the anti-PD-1 antibody nivolumab in ccRCC was obtained from the accompanying paper^64^. CM-025 was a randomized phase III trial which demonstrated an improved overall survival (OS) with nivolumab over the mTOR inhibitor everolimus. Patients were included based on the availability of RNAseq data and information on rhabdoid differentiation. After excluding cases with incomplete annotations, CD38 expression levels were plotted and analyzed in R.

#### Data integration by PPI network analysis

For comparison of proteomic and transcriptomic datasets we used the results from the differential expression and GO enrichment analyses performed for each data type independently between the normal and the three grades of the ccRCC. Clustering plots were generated using all genes with standard deviation in log2 fold change between samples of at least 0.5 and a maximum of 2000 genes. To highlight the synergies between the measured proteomics and transcriptomics profiles, we performed integrated network analysis using the NOODAI software platform^35^. Differentially expressed elements of the proteomics and transcriptomics analysis were used as input and searched against the default publicly available protein-protein interaction (PPI) databases containing only high confidence human interactors from STRING^82^, BioGRID^83^ and IntACT^84^ databases were collected. For STRING (version 11.5) complete interactome data from all sources was filtered to include only entries with a combined score above 0.7. For BioGRID (version 4.4.218), a mitab data file that contained a dataset of interactors with physical interactions supported by independent validations was used. It was filtered to exclude entries without an associated confidence value. For the IntACT database, the psimitab from 13/07/2022 was used and filtered to keep only interactions with a confidence score above 0.7. Interaction pairs obtained from the different databases were overlapped and merged into a joint dataset.

This was used in NOODAI to construct a network, in which only interaction pairs with both members significantly differentially expressed in at least one of the data layers (transcriptomics or proteomics) were retained. Only the network with the highest number of connected members was further analyzed (without allowing edge repetition). In order to identify highly connective central nodes in the networks, the default current flow betweenness centrality metric was used^85^.

Functional network modules^86^ were subsequently identified through NOODAI by using the MONET software^87^. Modularity optimization with undirected edges was selected, setting the target average node degree within the identified modules to 10. The modules were default sorted based on the number of nodes they contained. To assess the robustness of the network decomposition and to identify the main signaling axes present in our PPI networks, NOODAI performed pathways overrepresentation analysis for individual modules by searching against the pre-loaded Reactome and KEGG databases. By default, background datasets were composed of all genes and proteins that were used to construct the full-scale network of the respective conditions. Results were filtered to retain significant pathways with at least five significant hits and an FDR below 0.05 (Benjamini-Hochberg^88^ correction of Fisher’s exact test p-values)., Non-specific pathways with more than 300 members were omitted. Pathways that did not have more than two different overrepresented members were considered redundant and were investigated in the context of their explanatory power. For the visualization of results, NOODAI generated circular charts that highlighted the top 30% of most central nodes (based on the current-flow betweenness centrality). For selected modules and full-size networks, the graphical representation was done in Cytoscape 3.10.1^89^ using the webtool-generated edge files. For edge routing, the *organic routing* from yFiles plugin was used^90^. Additional R packages used for data handling and visualization of the NOODAI results included RColorBrewer^91^, readr^92^, stringr^93^, gtools^94^, gridBase^95^, ComplexHeatmap^96^, tidygraph^97^, biomaRt^98^ and readxl^99^.

## Author Contributions

H.A.B, P.H. and H.M. conceptualized the study. H.A.B. designed the experiments. H.A.B and N.J.R. selected and annotated clinical samples. E.M., A.K. and F.K. developed image analysis methodology and software. S.K. and H.A.B.and H.A.B. performed sample preparation, developed and performed sample preparation, developed and performed RNAseq analysis. D.R., S.P. and J.G. developed and performed mass-spectrometry analysis. C.L., P.L., N.Z. and T.T. conducted data analysis and integration. N.J.R. and H.M. analyzed histological data and provided scores of IHC marker expression. H.A.B, E.M., N.Z., T.T., N.J.R., M.B., H.M. and P.H. contributed to the interpretation of the results. H.A.B. wrote the original manuscript and all authors reviewed and edited it. H.A.B., P.H. and H.M. provided funding.

## Acknowledgement

From the Department of Pathology and Molecular Pathology at USZ, we acknowledge Susanne Dettwiler, Fabiola Prutek and Adriana von Teichmann for technical assistance and Prof. Dr. Peter Schraml for critical discussion. This work was supported by grants from the Edoardo R., Giovanni, Giuseppe und Chiarina Sassella-Foundation and the Julius Müller Foundation (to H.A.B.).

## Supplementary Figures

**Supplementary Figure 1:**
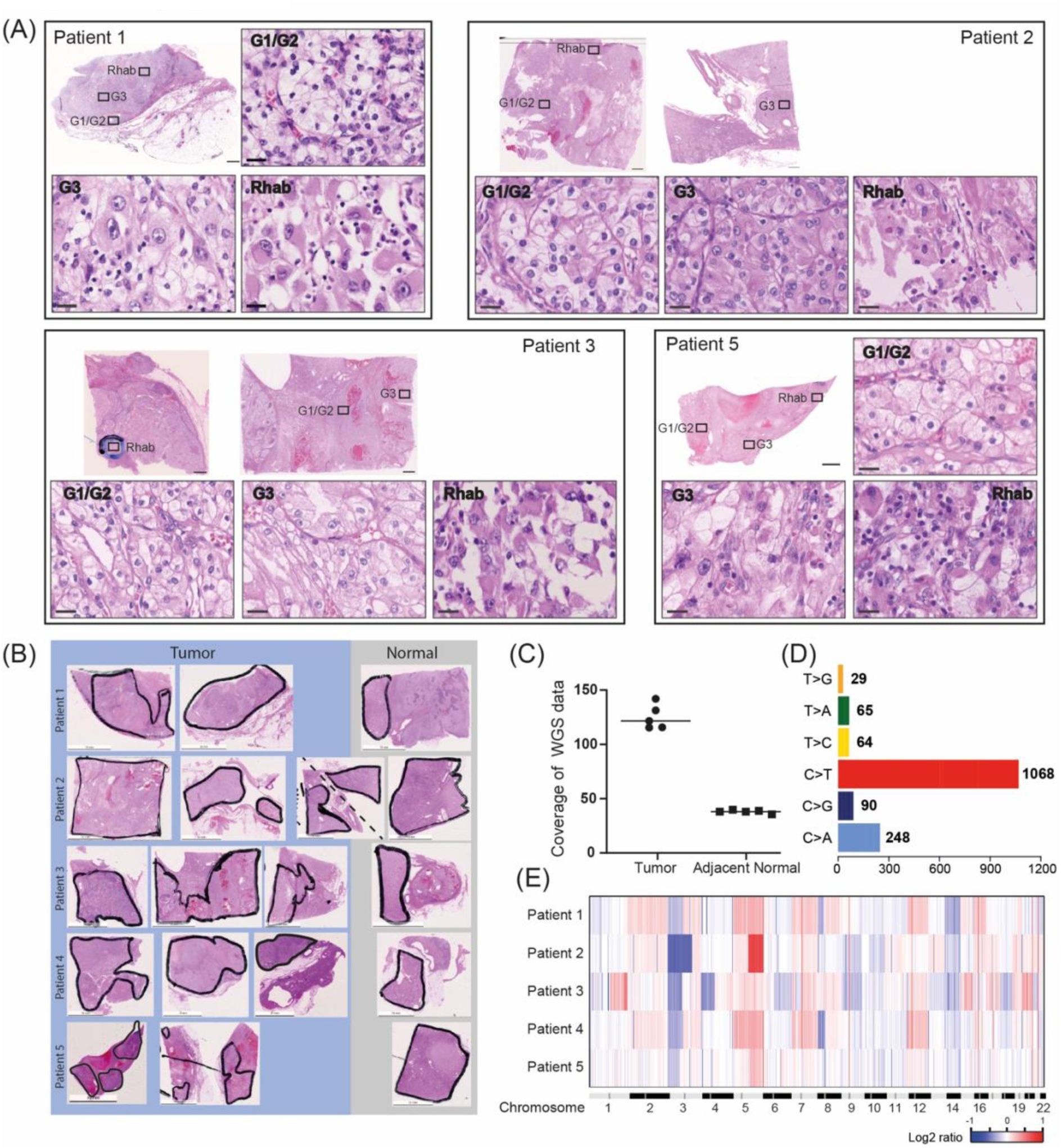
Image-guided single cell isolation and multi-omics analysis of ccRCC with pronounced morphological heterogeneity and rhabdoid de-differentiation. (A) Representative H&E images indicating ITH of grade features: Grade 1/2 (low-grade, G1/2), grade 3 (high-grade, G3) and grade 4 ccRCC with rhabdoid differentiation (Rhab). Scale bars: WSI – 2 mm, high magnification panels - 100 μm. (B) Overview images of WSI indicating the areas that were dissected to perform WGS. (C) Mean coverage of ccRCC and adjacent normal kidney in WGS data for all five cases. (D) Whole-genome mutation spectra indicating SNV classes, and their frequency, in our data highlighting the presence of C>T transitions caused by cytosine deamination from formalin fixation of the tissue samples. (E) CNV profiles determined from WGS data. CNV loss/gain are indicated by color coding, blue indicated loss and red indicated gain.

**Supplementary Figure 2:**
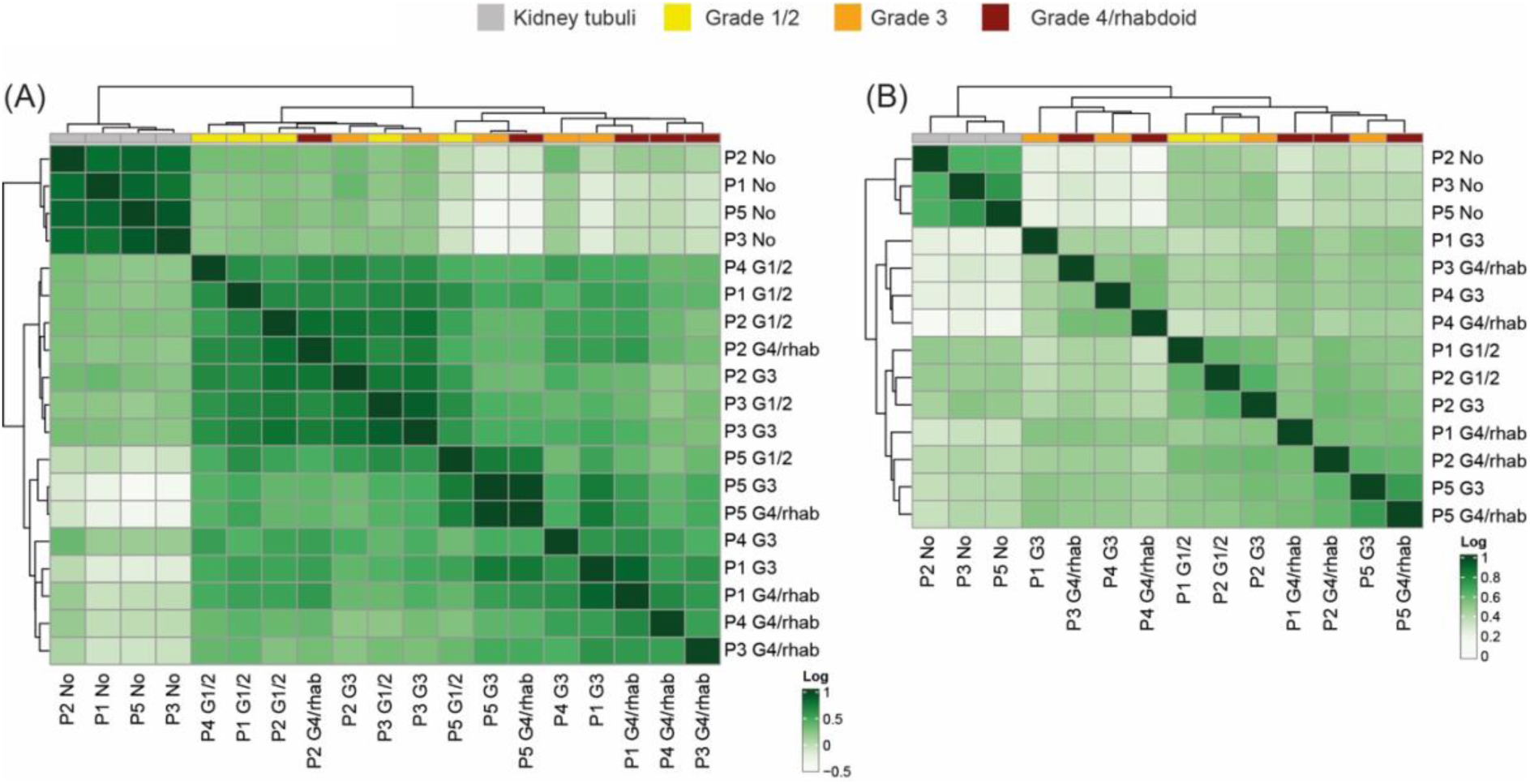
Analysis of mRNA and protein expression from individually selected cells with different grade morphologies. (A) Correlation analysis for top100 variance features of the transcriptomic dataset obtained from interrogating cells isolated from normal kidney tubuli tissue or ccRCC with distinct grade morphologies (N=19). Correlation coefficients are indicated by the color code. (B) Correlation analysis for proteomics samples isolated from normal kidney tubuli tissue or ccRCC with distinct grade morphologies (N=14). Correlation coefficients are indicated by the color code.

**Supplementary Figure 3:**
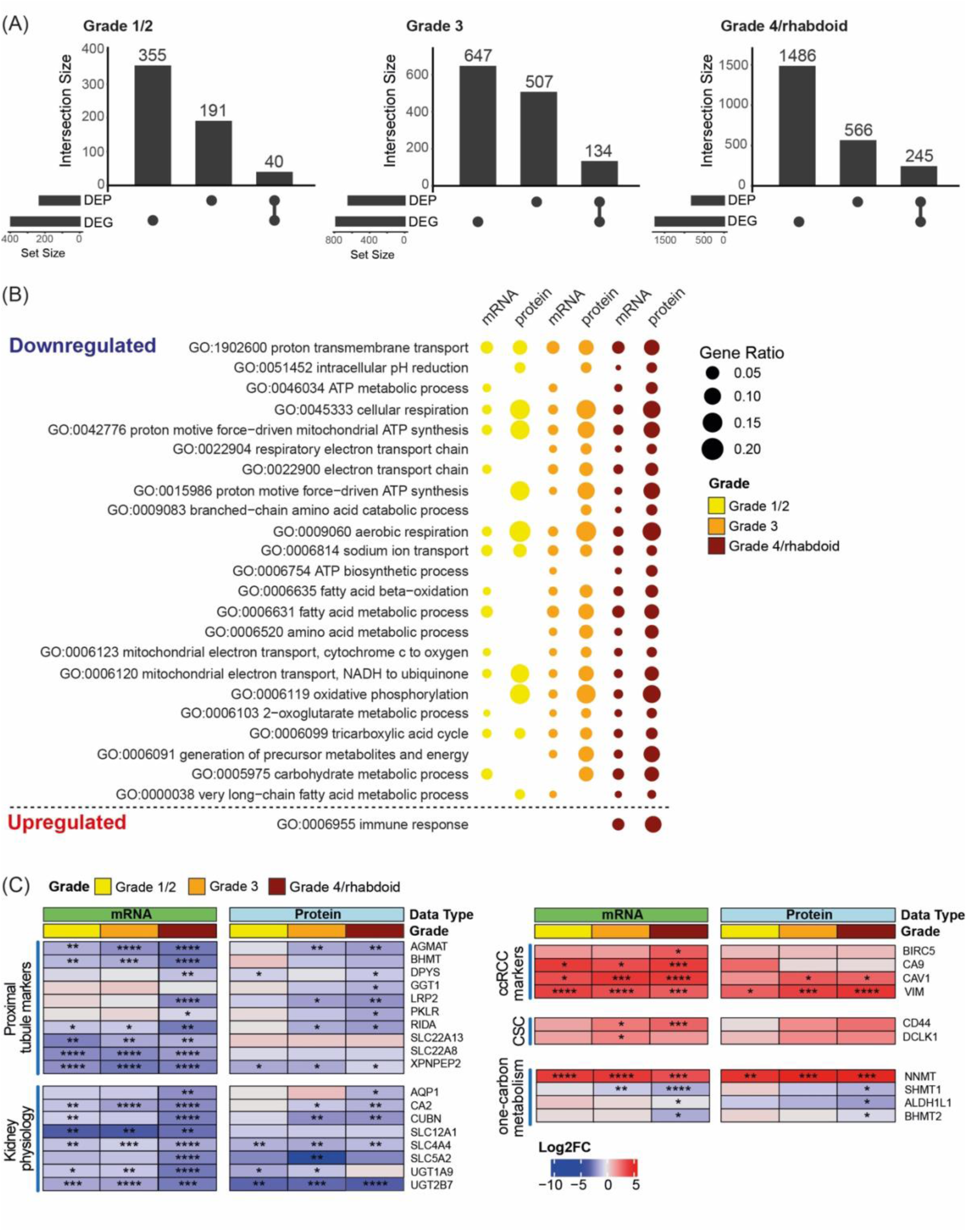
Functional overlap between mRNA and protein data. (A) Upset plots indicating the number of significants for mRNA and proteomics datasets for each grade group and their overlap. (B) Bubble plot depicting all 24 shared GO terms for mRNA and protein differential expression in the grade 4/rhabdoid class of ccRCC cells and their overlap with samples from other grade morphologies. (C) Heatmaps depicting selected alterations that mark kidney cell identity (proximal tubule cells markers and factors implicated in physiological kidney functions) as well as common ccRCC markers, cancer stem cell markers and factors implicated in one-carbon metabolism. Representative genes and proteins are depicted as log2FC with respect to adjacent normal cells. FDR is indicated by "****" <= 0.0001, "***" <= 0.001, "**"<= 0.01, "*" <= 0.05.

**Supplementary Figure 4:**
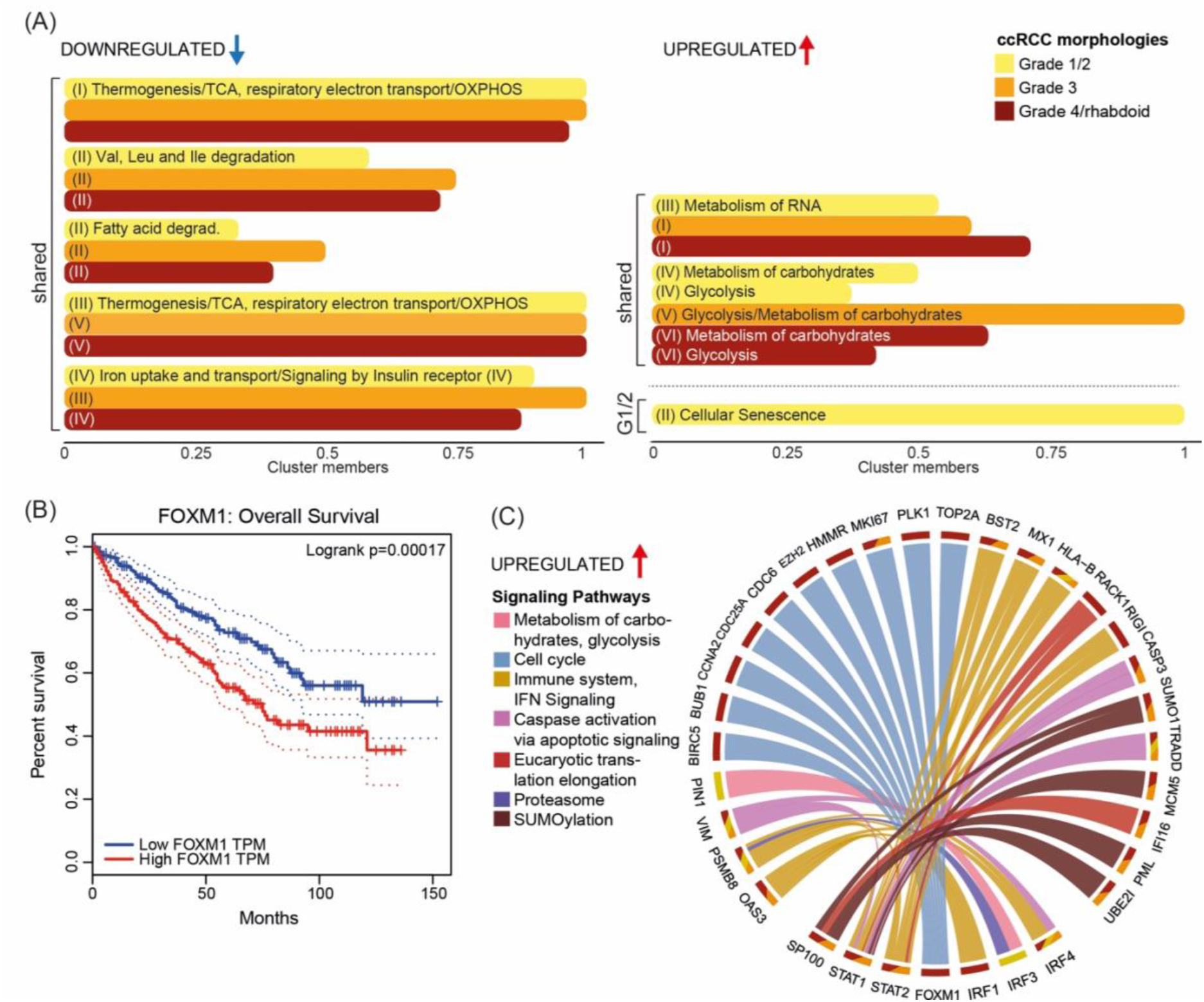
Multi-omics data integration uncovers distinct de-regulated processes in aggressive cells and their transcriptional regulation. (A) Biological processes enriched in modules shared by Grade 1/2 ccRCC grade morphology with at least 10 members following decomposition of the PPI networks in (A) based on KEGG and Reactome annotations. Enriched terms were determined by an FDR < 0.05 (Fisher’s exact test with Benjamini-Hochberg correction). Only non-redundant terms are presented, except where additional terms provided complementary information. The fraction of proteins in each module with the corresponding annotation is provided. (B) Kaplan-Meier survival analysis of FOXM1 in TCGA-KIRC dataset (n = 516) showing overall survival. HR (high) = 0.66 p(HR) = 0.0079. (C) Circular diagram highlighting the observed interaction between upregulated TFs (bottom) and their targets (top) within the top 30% nodes of the PPI networks based on the current-flow betweenness centrality for the three cell morphologies analyzed individually. Tracks mark the grade groups in which they were found to be significantly dysregulated (corresponds to color-code in (A)) and the bands color-code for the associated signaling pathways.

**Supplementary Figure 5:**
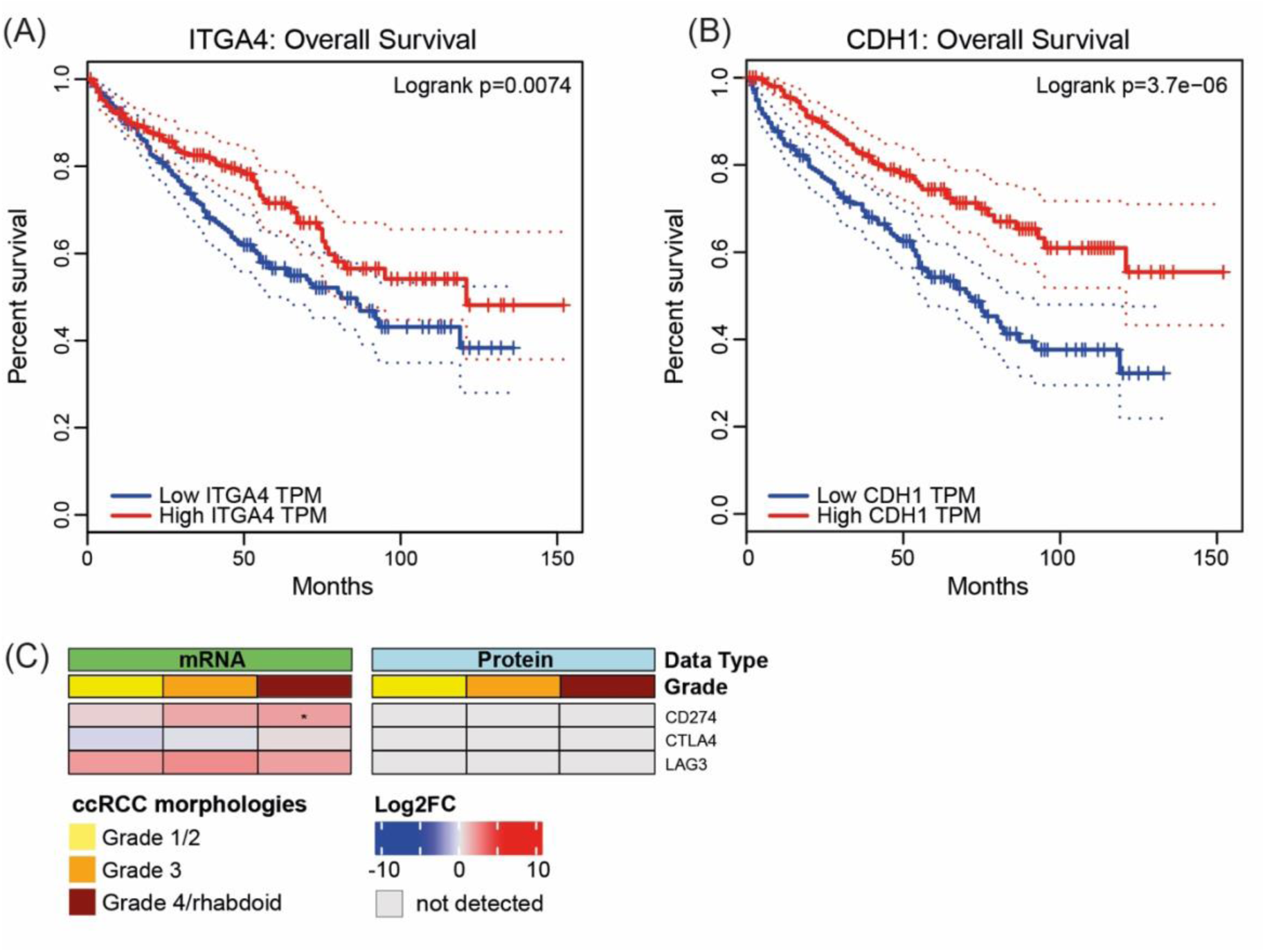
Grade 4/rhabdoid ccRCC cells show distinct expression profiles. (A) Kaplan-Meier survival analysis of ITGA4 with HR (high) = 1.8 p(HR) = 0.0022. (B) As in (A) for CDH1 with HR (high) = 0.48 p(HR) = 5.9e-06. (C) Heat maps highlighting selected elements associated with the immune system-related pathways and their respective dysregulation during ccRCC de-differentiation on both data layers. Alterations of representative genes and proteins are depicted as log2FC with respect to adjacent normal cells. FDR is indicated by"****" <= 0.0001, "***" <= 0.001, "**"<= 0.01, "*" <= 0.05.

